## Supplementary methods and data for "The place-cell representation of volumetric space in rats"

### Supplementary information

Supplementary data (15 figures, 1 table, 2 videos)

Supplementary methods (4 figures, 1 table, 2 videos)

All figures are available for download in high resolution format at: DOI: 10.5522/04/9977435

### Supplementary data

*Movement patterns in the lattice mazes*

As reported in the main text, rats explored both lattice mazes fully but explored the tilted lattice less ($\chi^2(1) = 19.4$, $p < .0001$, $\eta_p^2 = 0.34$, K-W) although the small number of tilted maze sessions (N = 16) should be taken into consideration when interpreting this result (Fig. S1a-b). In both configurations there was a differential distribution of time spent in the vertical dimension: rats spent a median of 1.63 times longer in the bottom half of the maze in its aligned configuration (Z = 2.31, $p = .021$, U3 = 0.11, WSR) and 1.95 times in the tilted configuration (Z = 1.46, $p = .14$, U3 = 0.25, WSR): these did not differ ($\chi^2(1) = 0.095$, $p = .76$, $\eta_p^2 = 0.007$, K-W, Fig. S1c&g). Rats also tended to remain close to maze boundaries (median time outer half volume / inner half volume: arena: 2.60, Z = 3.18, $p = .001$, U3 = 0, lattice: 1.77, Z = 2.67, $p = .008$, U3 = 0, tilted: 1.73, Z = 1.46, $p = .144$, U3 = 0.25, WSR tests) and to a similar extent in all three ($\chi^2(2) = 3.15$, $p = .21$, $\eta_p^2 = 0.12$, K-W, Fig. S1d&g). This is interesting because the lattice mazes had no solid boundaries, the boundaries were defined solely by the termination of maze struts.

In the open field arena animals unsurprisingly moved significantly slower along the Z-axis while the X and Y axes did not differ (median speed in X,Y & Z: 0.081, 0.081 & 0.025 m/s, $\chi^2(2) = 19.8$, $p = .00005$, $\eta_p^2 = 0.51$, FT, post-hoc: X vs Z & Y vs Z, $p < .002$, X vs Y, $p > .99$, Fig. S1e). In the aligned lattice movements along the Z-axis were significantly slower (median speed in X, Y & Z: 0.055, 0.054 & 0.032 m/s, $\chi^2(2) = 14.0$, $p = .0009$, $\eta_p^2 = 0.52$, FT, post-hoc: X vs Z & Y vs Z, $p < .02$, X vs Y, $p > .99$, Fig. S1e). In the tilted lattice animals moved slightly faster along the Z-axis (median speed in X,Y & Z: 0.041, 0.043 & 0.049 cm/s, $\chi^2(2) = 8.0$, $p = .018$, $\eta_p^2 = 0.67$, FT, post-hoc: X vs Y & Y vs Z, $p > .47$, X vs Z, $p = .014$) but they moved at an equivalent speed along the (now rotated) maze axes, which we labelled A, B and C (median speed in A,B & C: 0.042, 0.042 & 0.041 m/s, $\chi^2(2) = 2.0$, $p = .37$, $\eta_p^2 =$

0.17, FT, Fig. S1e). The small number of tilted lattice sessions (N = 16) should be taken into consideration when interpreting these results.

Animals spent significantly less time moving at slower speeds (F(24,1225) = 126.0, $p$ < .0001, $\eta_p^2$ = 0.71) but this did not differ significantly between environments (F(2,1225) = 0.2, $p$ = .82, $\eta_p^2$ < 0.001) nor was there a significant interaction between the two (F(48,1225) = 1.2, $p$ = .17, $\eta_p^2$ = 0.045, Univariate ANOVA comparing effects of speed and environment on dwell time, Fig. S1f).

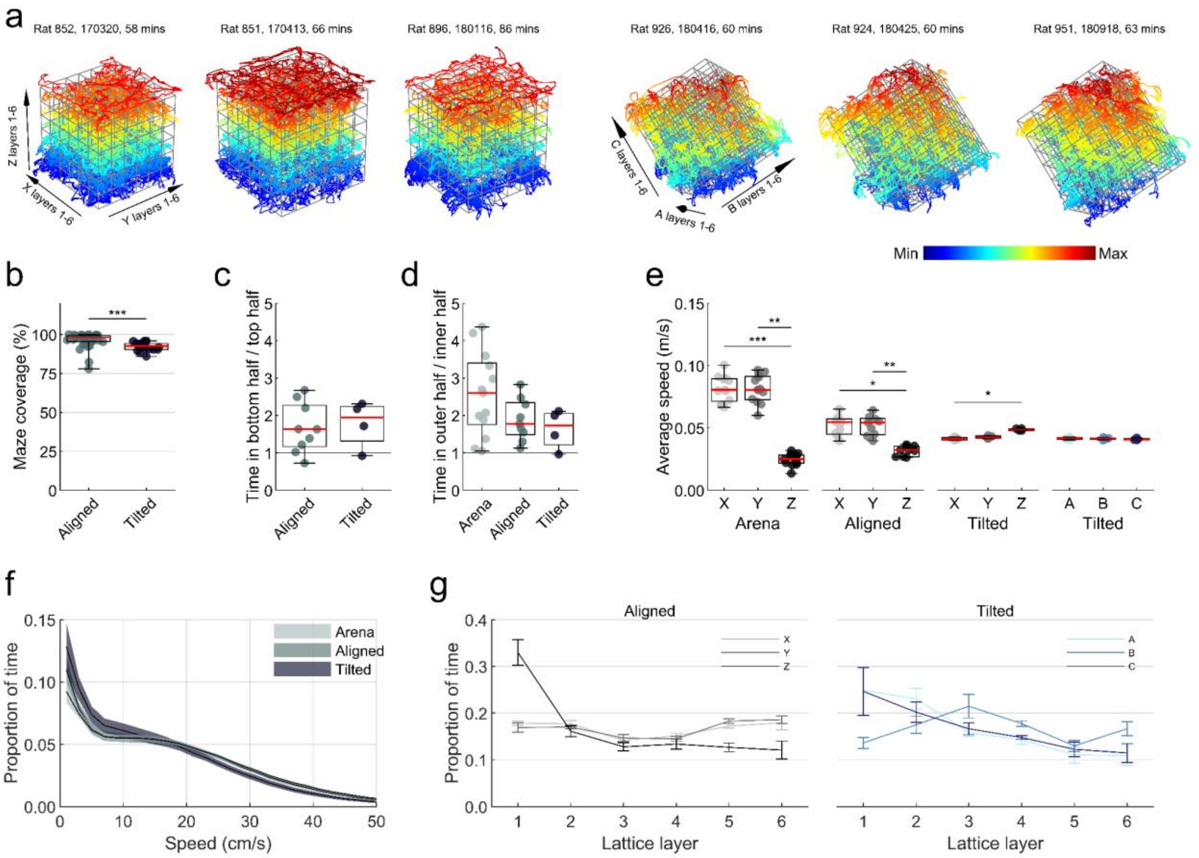


**Fig. S1.** Movement patterns in the lattice mazes. **a** Position tracking from 3 representative example aligned lattice sessions (left) and 3 tilted lattice sessions (right). Color denotes vertical height. Animals explored the full extent of the lattice, although some would not explore the very top level of the aligned lattice (first example) and some would spend more time in the bottom level (third example). Often animals would not explore the topmost vertex of the tilted lattice (fourth example). **b** Markers represent sessions. The percentage coverage (i.e. visited lattice nodes) in each maze. **c** Markers represent animals. The average ratio of time spent in the bottom half of each lattice compared to the top. **d** Markers represent animals. The average ratio of time spent in the outer half of each lattice compared to the inner half. **e** Markers represent animals. The average speed at which rats moved along each maze axis. **f** The mean and SEM proportion of time animals spent moving at various running speeds, averaged across sessions. **g** Left; aligned lattice, right; tilted lattice. Time spent in each lattice layer (see **a**), averaged across animals. Closer inspection of the distribution of dwell time in the mazes reveals that animals spent more time in the outer layers of the lattice mazes, especially in the bottom layer of the aligned lattice and the layers of the tilted lattice closest to the floor. Source data are provided as a Source Data file.

*Summary statistics of units and place cells*

**Table S1**

*Summary statistics of units and place cells recorded in all 3 environments*

| Maze | Rat | Sessions | | Cells | | Place cells | | Place fields | | Fields per Cell | Field elongation | | Spatial information | |
|---|---|---|---|---|---|---|---|---|---|---|---|---|---|---|
| | | n | % of total | n | % of total | n | % of total | n | % of total | | μ | s.d. | μ | s.d. |
| | 750 | 1 | 2.0 | 11 | 1.0 | 1 | 0.2 | 1 | 0.1 | 1.00 | 1.27 | 0.00 | 1.10 | 1.46 |
| | 770 | 2 | 4.0 | 25 | 2.4 | 21 | 3.3 | 23 | 3.3 | 1.10 | 2.01 | 0.70 | 1.16 | 0.76 |
| | 775 | 4 | 8.0 | 96 | 9.0 | 60 | 9.6 | 50 | 7.2 | 0.83 | 1.89 | 0.83 | 1.48 | 1.36 |
| | 850 | 2 | 4.0 | 24 | 2.3 | 6 | 1.0 | 9 | 1.3 | 1.50 | 1.72 | 0.64 | 1.38 | 1.44 |
| | 851 | 5 | 10.0 | 62 | 5.8 | 16 | 2.6 | 24 | 3.5 | 1.50 | 1.69 | 0.46 | 1.38 | 1.24 |
| | 852 | 14 | 28.0 | 186 | 17.5 | 75 | 11.9 | 85 | 12.3 | 1.13 | 1.72 | 0.49 | 1.60 | 1.41 |
| Arena | 853 | 3 | 6.0 | 45 | 4.2 | 20 | 3.2 | 17 | 2.5 | 0.85 | 1.84 | 0.66 | 1.30 | 1.11 |
| | 894 | 3 | 6.0 | 67 | 6.3 | 37 | 5.9 | 37 | 5.3 | 1.00 | 1.56 | 0.45 | 0.88 | 0.72 |
| | 896 | 5 | 10.0 | 196 | 18.4 | 163 | 26.0 | 179 | 25.8 | 1.10 | 1.85 | 0.59 | 1.40 | 1.01 |
| | 923 | 2 | 4.0 | 9 | 0.9 | 3 | 0.5 | 4 | 0.6 | 1.33 | 1.88 | 0.80 | 0.58 | 0.66 |
| | 924 | 2 | 4.0 | 5 | 0.5 | 3 | 0.5 | 3 | 0.4 | 1.00 | 1.40 | 0.23 | 1.16 | 0.71 |
| | 926 | 9 | 18.0 | 295 | 27.7 | 192 | 30.6 | 219 | 31.6 | 1.14 | 1.85 | 0.66 | 1.55 | 1.34 |
| | 951 | 5 | 10.0 | 43 | 4.0 | 31 | 4.9 | 42 | 6.1 | 1.35 | 1.83 | 0.67 | 0.98 | 0.62 |
| Total | 13 | 50 | - | 1064 | - | 628 | - | 693 | - | - | - | - | - | - |
| | 750 | 1 | 2.9 | 11 | 1.5 | 3 | 0.6 | 2 | 0.3 | 0.67 | 1.70 | 0.31 | 1.03 | 1.19 |
| | 770 | 2 | 5.9 | 25 | 3.5 | 18 | 4.2 | 26 | 4.3 | 1.44 | 1.64 | 0.39 | 1.88 | 2.39 |
| | 775 | 4 | 11.8 | 96 | 13.5 | 60 | 13.9 | 73 | 12.1 | 1.22 | 1.60 | 0.43 | 1.75 | 1.82 |
| | 850 | 2 | 5.9 | 24 | 3.4 | 17 | 3.9 | 26 | 4.3 | 1.53 | 1.81 | 0.61 | 1.48 | 1.04 |
| Aligned | 851 | 5 | 14.7 | 62 | 8.7 | 22 | 5.1 | 27 | 4.5 | 1.23 | 1.66 | 0.77 | 1.26 | 1.35 |
| | 852 | 14 | 41.2 | 186 | 26.1 | 80 | 18.5 | 100 | 16.6 | 1.25 | 1.61 | 0.44 | 1.47 | 1.32 |
| | 853 | 3 | 8.8 | 45 | 6.3 | 18 | 4.2 | 28 | 4.6 | 1.56 | 1.65 | 0.43 | 1.92 | 1.62 |
| | 894 | 3 | 8.8 | 67 | 9.4 | 40 | 9.3 | 63 | 10.4 | 1.57 | 1.75 | 0.59 | 0.73 | 0.70 |
| | 896 | 5 | 14.7 | 196 | 27.5 | 174 | 40.3 | 259 | 42.9 | 1.49 | 1.71 | 0.49 | 1.31 | 0.97 |
| Total | 9 | 34 | - | 712 | - | 432 | - | 604 | - | - | - | - | - | - |
| | 923 | 2 | 12.5 | 9 | 2.6 | 4 | 1.48 | 10 | 2.5 | 2.50 | 1.75 | 0.36 | 1.25 | 1.03 |
| Tilted | 924 | 2 | 12.5 | 5 | 1.4 | 3 | 1.11 | 2 | 0.5 | 0.67 | 1.39 | 0.14 | 1.43 | 0.90 |
| | 926 | 9 | 56.3 | 295 | 83.8 | 232 | 85.6 | 335 | 84.2 | 1.44 | 1.69 | 0.51 | 1.40 | 1.17 |
| | 951 | 5 | 31.3 | 43 | 12.2 | 32 | 11.8 | 51 | 12.8 | 1.59 | 1.68 | 0.50 | 1.14 | 0.76 |
| Total | 4 | 16 | - | 352 | - | 271 | - | 398 | - | - | - | - | - | - |

*Example place cells, aligned lattice*

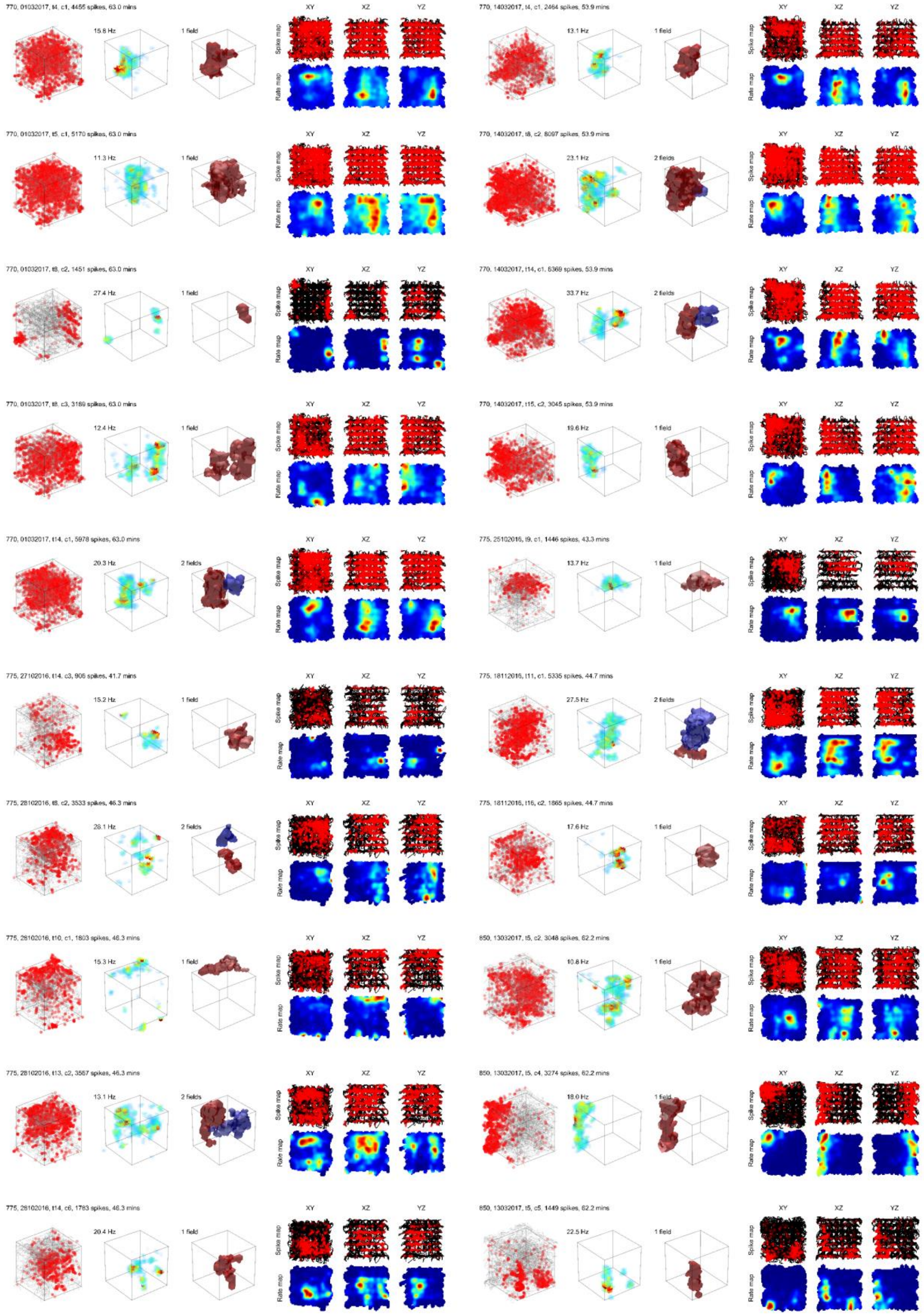

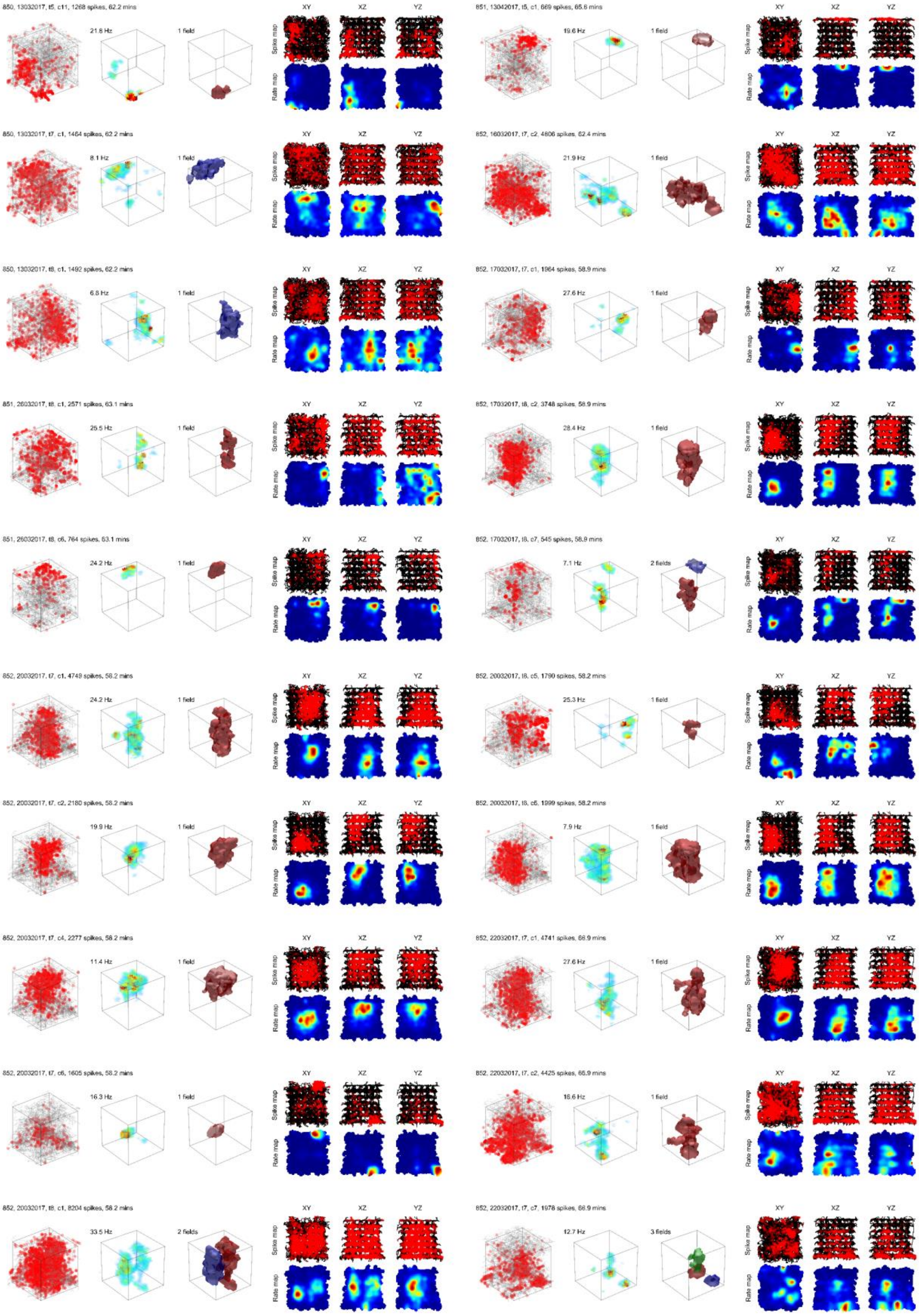
XY
XZ
YZ
Spike map
Rate map

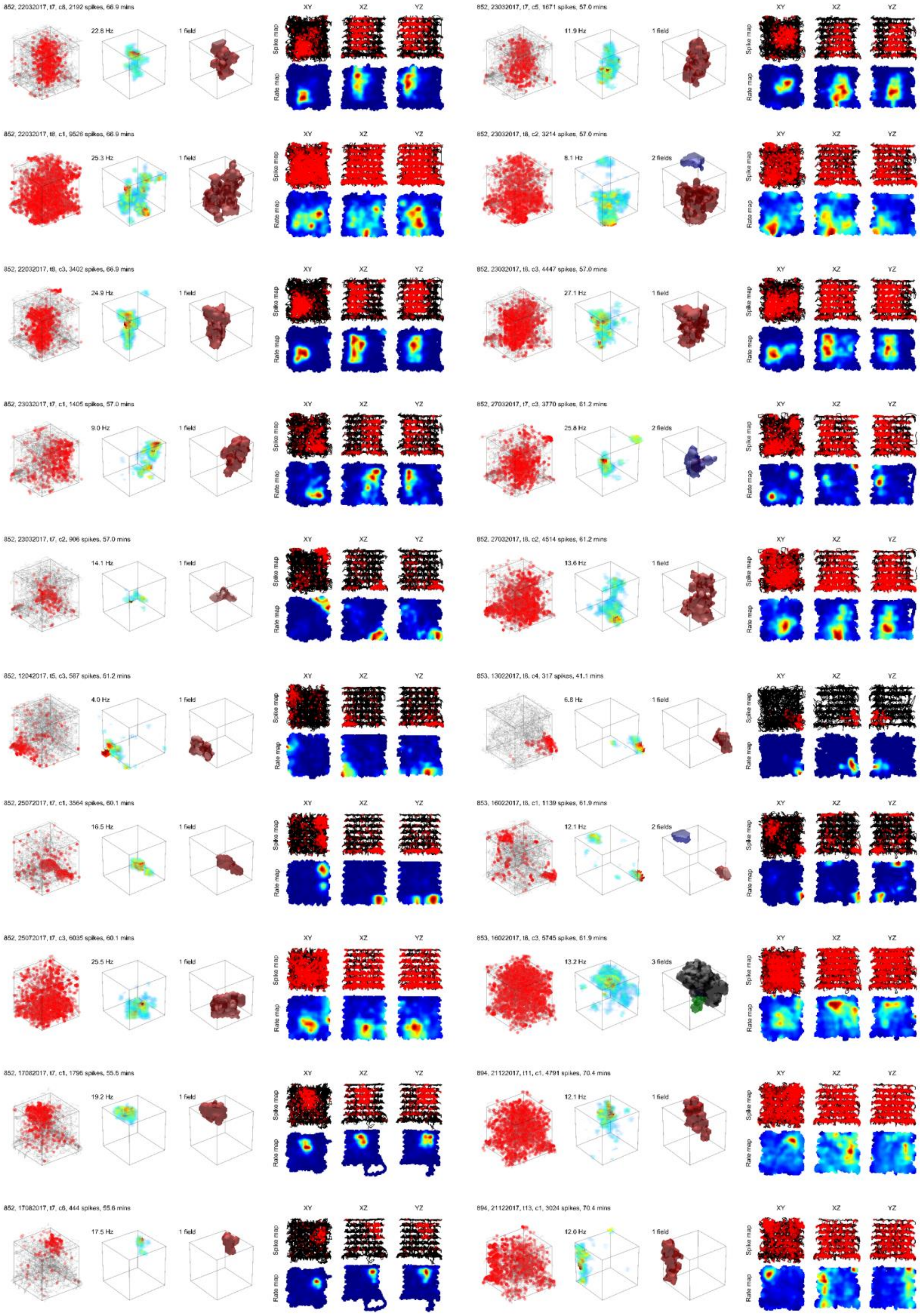

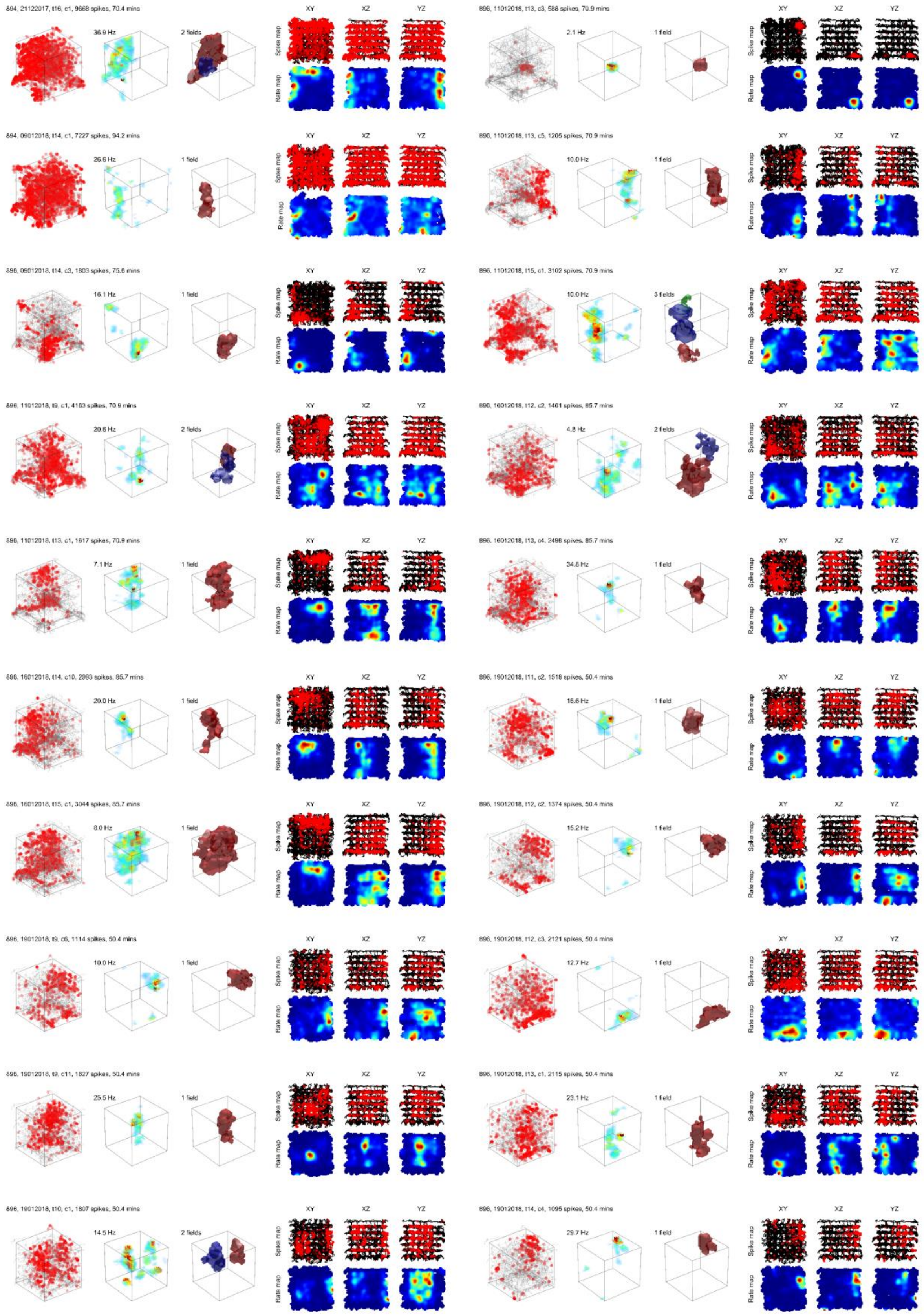

**Fig. S2:** Preceding 4 pages; activity of representative place cells recorded in the aligned lattice maze. For rotating plots see Supplementary Video S1. These cells were chosen to reflect the varied firing patterns observed in the lattice maze, across as many sessions and animals as possible, but were also chosen for their simpler firing patterns which are easier to visualize here in 2D. Two cells are shown per row. Leftmost plots show the animal's path (grey lines) and the cell's spikes are represented by red markers. A volumetric firing rate map of the data is shown to the right of the spike plot. High firing rates are represented by hot colors, low firing rates are represented by cold colors. Voxels with low firing rates are more transparent. Additionally, unvisited voxels and voxels containing firing rate values <10% of the map maximum are completely transparent. To the right of the firing rate map is a volumetric outline of detected place fields, represented by colored polygons. Different colors represent different fields. The smaller plots to the right of this show the data when projected onto the cardinal planes. The top row shows the projected spike and position data, the bottom row shows the firing rate map produced using these projected data. The rat number, recording date, tetrode and cluster of each cell is shown above the spike and position plot.

*Example place cells, tilted lattice*

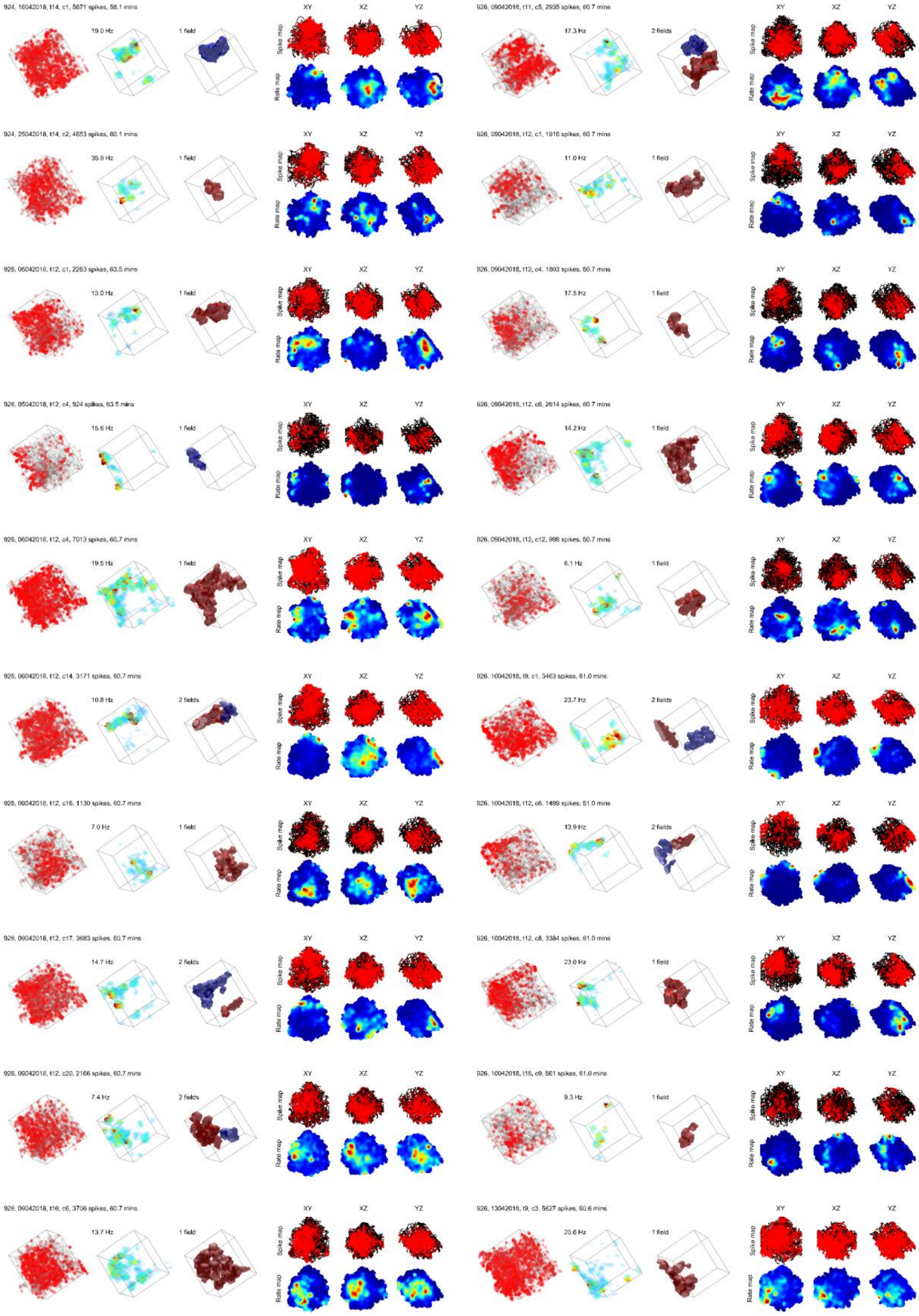

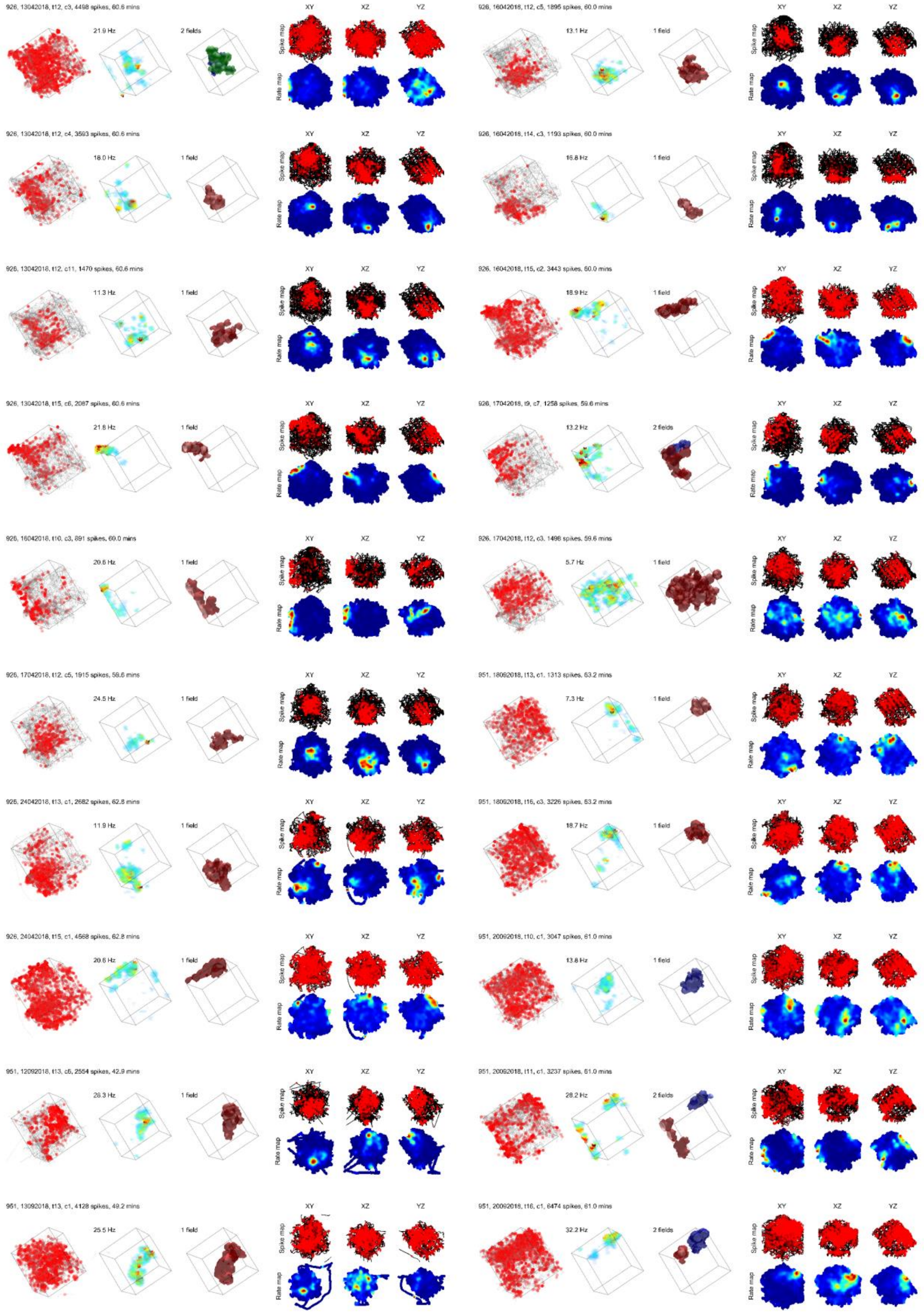

926, 13042018, t12, c3, 4498 spikes, 60.6 mins
21.9 Hz
2 fields
926, 13042018, t12, c4, 3593 spikes, 60.6 mins
18.0 Hz
1 field
926, 13042018, t12, c11, 1470 spikes, 60.6 mins
11.3 Hz
1 field
926, 13042018, t15, c6, 2087 spikes, 60.6 mins
21.8 Hz
1 field
926, 16042018, t10, c3, 891 spikes, 60.0 mins
20.6 Hz
1 field
926, 17042018, t12, c5, 1915 spikes, 59.6 mins
24.5 Hz
1 field
926, 24042018, t13, c1, 2682 spikes, 62.8 mins
11.9 Hz
1 field
926, 24042018, t15, c1, 4568 spikes, 62.8 mins
20.6 Hz
1 field
951, 12092018, t13, c5, 2554 spikes, 42.9 mins
28.3 Hz
1 field
951, 13092018, t13, c1, 4128 spikes, 49.2 mins
25.5 Hz
1 field
926, 16042018, t12, c5, 1895 spikes, 60.0 mins
13.1 Hz
1 field
926, 16042018, t14, c3, 1193 spikes, 60.0 mins
16.8 Hz
1 field
926, 16042018, t15, c2, 3443 spikes, 60.0 mins
18.9 Hz
1 field
926, 17042018, t9, c7, 1258 spikes, 59.6 mins
13.2 Hz
2 fields
926, 17042018, t12, c3, 1498 spikes, 59.6 mins
5.7 Hz
1 field
951, 18092018, t13, c1, 1313 spikes, 63.2 mins
7.3 Hz
1 field
951, 18092018, t16, c3, 3226 spikes, 63.2 mins
18.7 Hz
1 field
951, 20092018, t10, c1, 3047 spikes, 61.0 mins
13.8 Hz
1 field
951, 20092018, t11, c1, 3237 spikes, 61.0 mins
28.2 Hz
2 fields
951, 20092018, t16, c1, 6474 spikes, 61.0 mins
32.2 Hz
2 fields

**Fig. S3:** Preceding 2 pages; activity of representative place cells in the tilted lattice maze, as in Fig. S2. For rotating plots see Supplementary Video S2.

*Recording stability*

We compared the firing activity between the first and second arena sessions as a measure of recording stability (Fig. S4a, Supp. Methods: *Recording stability*). When data were projected onto the XY axis, correlations between arena sessions were significantly higher than would be expected by chance (blue and red areas respectively, Fig. S4b). On top of this, the median observed correlation exceeded the 95$^{th}$ percentile of the shuffle distribution (blue and red vertical lines respectively, Fig. S4b). The same effects were observed when data were projected onto the XZ plane and when correlating whole, volumetric firing rate maps (Fig. S4b). However, although YZ projections were significantly higher than chance, their median did not exceed the 95$^{th}$ percentile.

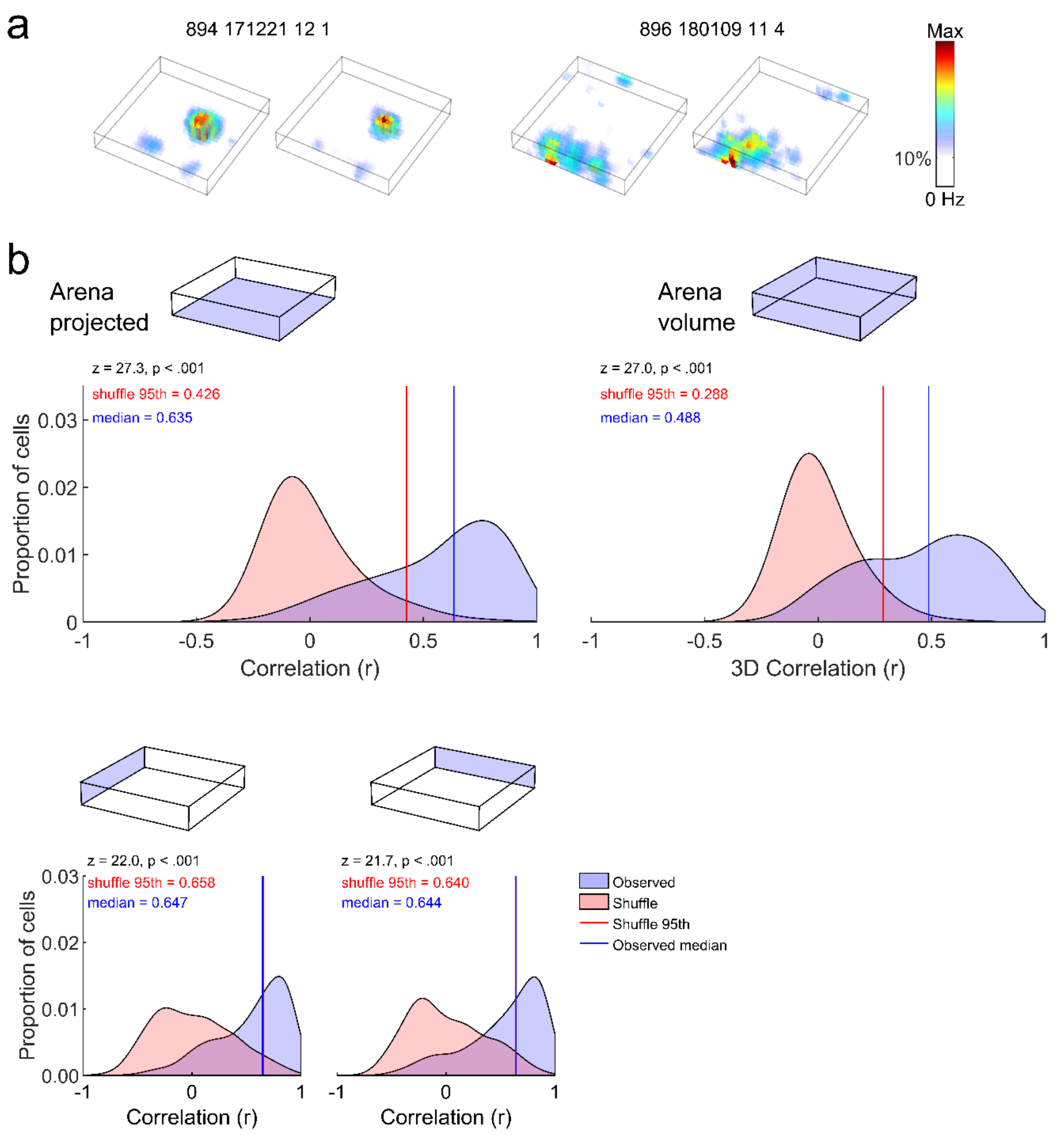


**Figure S4:** Stability of recording between first and second arena. **a** Consecutive open field arena volumetric ratemaps for two example cells. **b** Schematics indicate the type of comparison, either showing the plane of projection or highlighting the whole volume. Plots give the distribution of observed intra-trial correlation scores between the first and second open field sessions (blue shaded area) and between 1000 randomly shuffled cells (red shaded area). Black text gives the result of a right-tailed Wilcoxon rank sum test comparing the distributions. Red lines denote the 95th percentile rank position in the shuffled distribution, blue lines denote the median position in the observed distribution. The top left plot shows the result of this analysis when comparing the 1st and 2nd arena after projecting all data onto the XY plane. Smaller plots below this show the same when data are projected onto the XZ and YZ planes. Top right plot shows the result of this analysis when whole volumetric firing rate maps are compared without projection. Source data are provided as a Source Data file.

*Zingg shape categorisation*

Place fields took on different shapes in the mazes, most fields were elongated in the lattice mazes while they exhibited a flattened shape in the arena (Fig. S5). Significance was determined by a random shuffle of all place field axis lengths (Supp. Methods: *Zingg shape categorization*). In the arena more fields were bladed and oblate that would be expected by chance (36.8% and 26.6%, shuffle 99$^{th}$ percentiles: 22.1% and 24.8%). In the lattice maze fields were most often prolate (33.6%) but no field type exceeded chance. In the tilted lattice more fields were prolate than would be expected by chance (37.7%, shuffle 99$^{th}$ percentile: 35.0%).

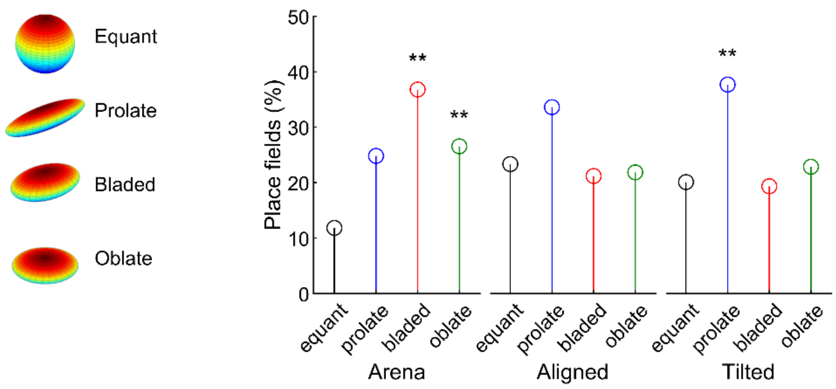


**Fig. S5:** Zingg shape classification of place fields in each maze environment. Equant fields are equidimensional, prolate fields resemble a cigar, oblate fields resemble a pancake and bladed fields are different along every dimension. Observed values were compared to a shuffle, asterisks indicate the value exceeded the 99$^{th}$ percentile of the shuffle. Source data are provided as a Source Data file.

*Trajectory downsampling*

To test whether the animals' biased movements in the aligned lattice affected our spatial information and correlational analyses we performed a downsampling procedure to correct for these effects (Fig. S6a, Supp. Methods: *Trajectory downsampling*). As reported in the main text, the spatial information exhibited by place cells in the aligned lattice differed along the X, Y and Z axes with horizontal slices along the Z-axis demonstrating the greatest spatial information (Fig. 8). In the downsampled data this effect was preserved (Fig. S6b). As reported in the main text, autocorrelation values were higher at longer distances in the aligned lattice along the Z-axis. This effect was also preserved after downsampling (Fig. S6c).

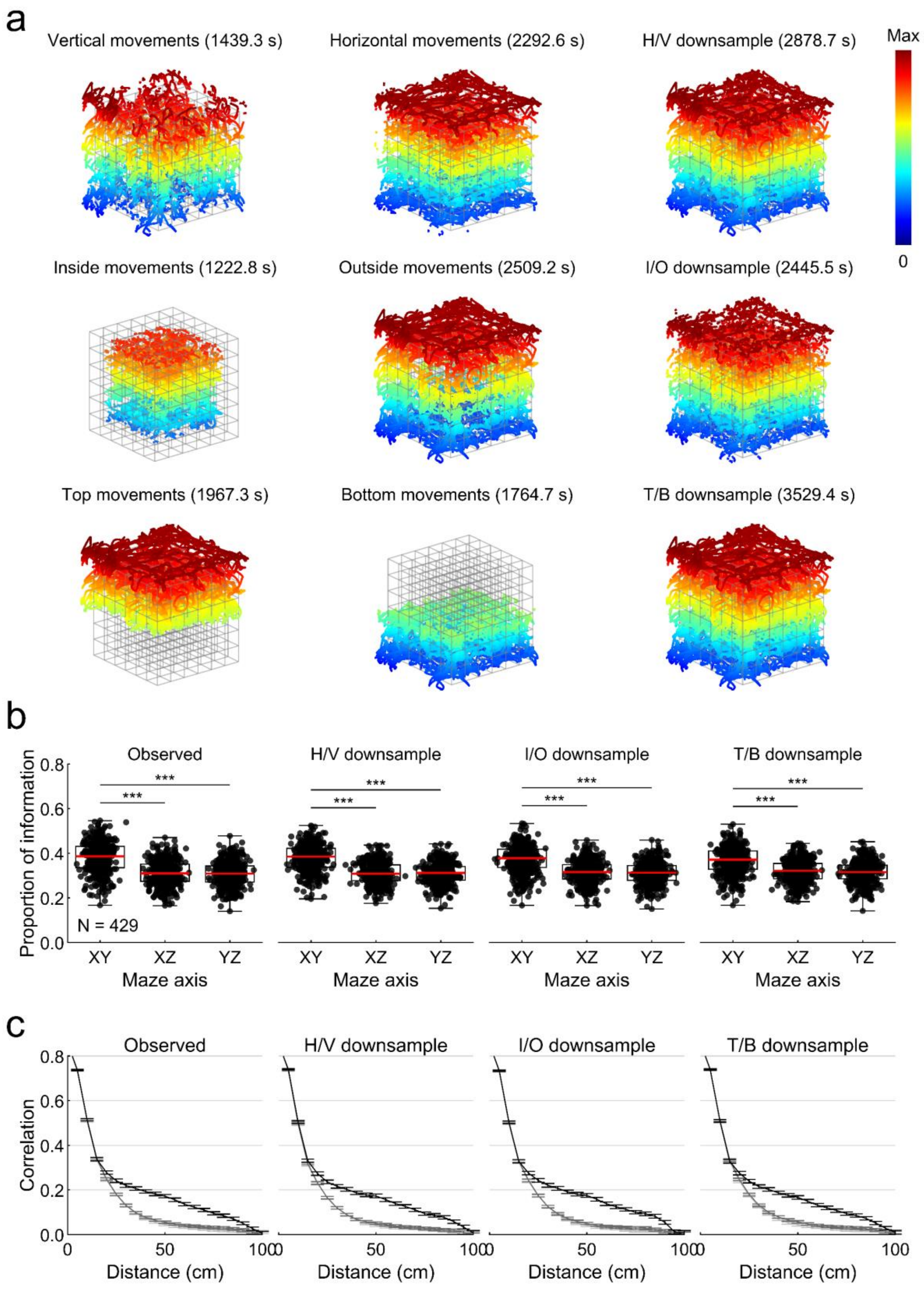

a
Vertical movements (1439.3 s)
Horizontal movements (2292.6 s)
H/V downsample (2878.7 s)
Max
0
Inside movements (1222.8 s)
Outside movements (2509.2 s)
I/O downsample (2445.5 s)
Top movements (1967.3 s)
Bottom movements (1764.7 s)
T/B downsample (3529.4 s)
b
Observed
H/V downsample
I/O downsample
T/B downsample
Proportion of information
N = 429
XY
XZ
YZ
Maze axis
c
Correlation
Distance (cm)


173

**Fig. S6:** The effect of downsampling trajectories on self-similarity and spatial information. **a** example trajectory of a rat recorded in the aligned lattice, filtered to include different subsets of data. Top row; the trajectory filtered to include only the periods where he was moving vertically (left), moving horizontally (middle), or a combination of these so that 50% of the data are horizontal and vertical movements (right). Middle row; same, but the data are filtered to include only data within the inner 50% volume of the lattice, the outer 50% volume or an equal combination of these. Bottom row, the same but for the top 50% of the lattice and the bottom 50% of the lattice. **b** The same spatial information analysis reported in the main text carried out on observed/unfiltered data (left) and on the 50% combinations (right plots). **c** The same autocorrelation analysis reported in the main text carried out on observed/unfiltered data (left) and on the 50% combinations (right plots). In all cases the effects remain the same in all forms of downsampled data. Source data are provided as a Source Data file.

*Field elongation*

In the aligned lattice the lengths of fields along X and Y were unimodal (deviation from unimodality: $p$ = .44, $p$ = .41 respectively) but the height of fields along the Z axis deviated from a unimodal distribution (deviation from unimodality: $p$ < .001) and instead formed a bimodal one (deviation from bimodality: $p$ = .46, all tests bootstrap modality tests). The left-hand peak of this bimodal distribution contains many horizontally elongated fields, while the smaller right-hand peak contains many vertically elongated fields, as would be expected (Fig. S7a). There was no significant deviation from modality for any axis in the tilted lattice ($p$ > .4 in all cases).

Place field elongation in the lattice mazes was weakly but significantly positively correlated with field centroid distance from maze center (Fig. S7b). There was no significant relationship between field elongation and experience in the mazes, although our animals were pre-exposed to small lattice mazes for weeks before recording (Fig. S7c, Methods: *Animals*). There was also no relationship between cluster quality measures and place field elongation ($L_{ratio}$: $r$ = 0.084, $p$ = .11; Iso-D: $r$ = -0.010, $p$ = .85; RPVs: $r$ = -0.100, $p$ = .06; spike amplitude: $r$ = -0.046, $p$ = .38; SNR: $r$ = -0.015, $p$ = .781, pairwise Spearman's correlations, Fig. S7d).

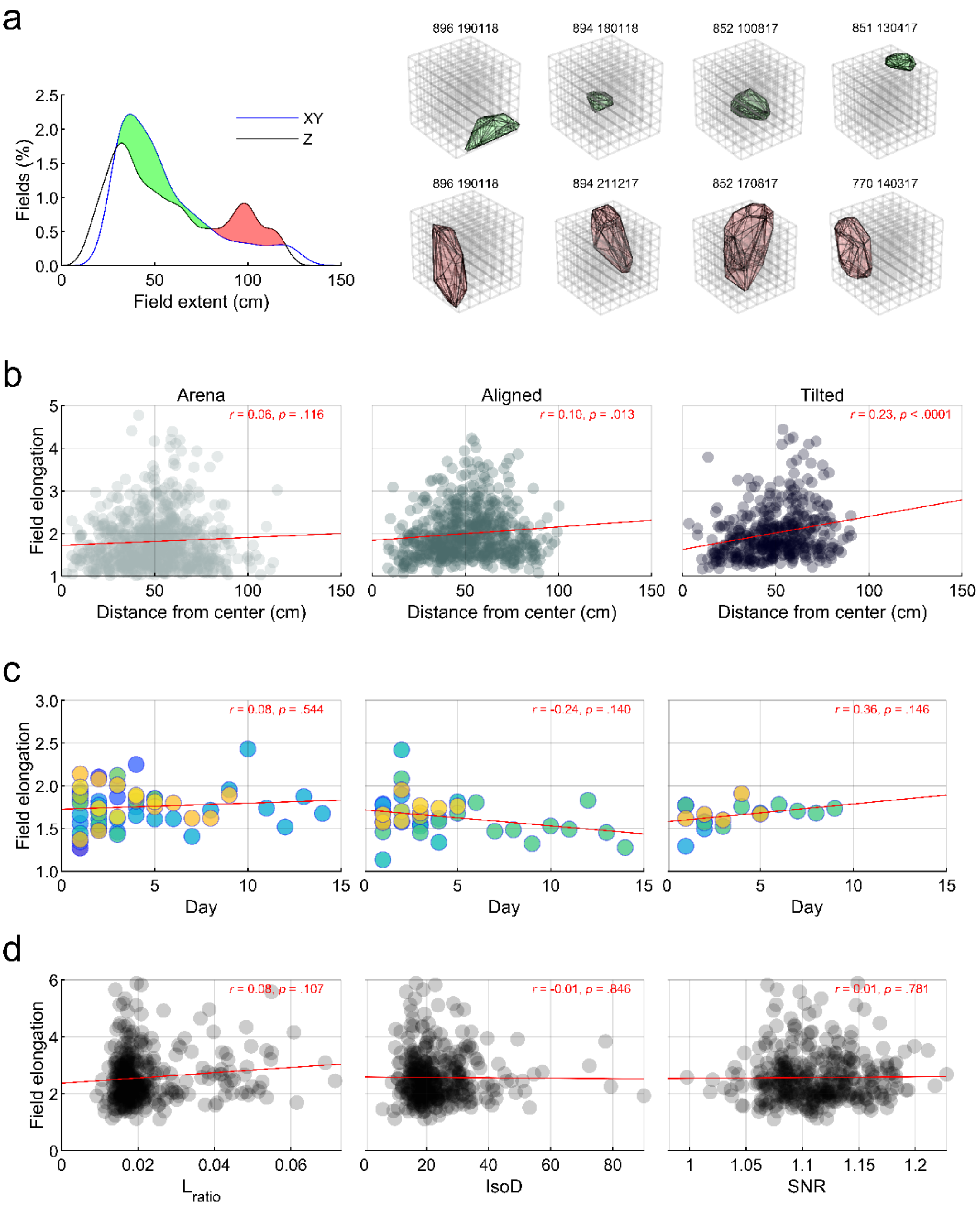
a
896 190118
894 180118
852 100817
851 130417
896 190118
894 211217
852 170817
770 140317
XY
Z
Fields (%)
Field extent (cm)
b
Arena
Aligned
Tilted
r = 0.06, p = .116
r = 0.10, p = .013
r = 0.23, p < .0001
Field elongation
Distance from center (cm)
c
r = 0.08, p = .544
r = -0.24, p = .140
r = 0.36, p = .146
Field elongation
Day
d
r = 0.08, p = .107
r = -0.01, p = .846
r = 0.01, p = .781
Field elongation
L ratio
IsoD
SNR

**Fig. S7:** Place field elongation, continued from Fig. 6. **a** Two distributions are shown, the average of the X & Y distributions in Fig. 6 (blue line) and the Z distribution. The regions between these curves correspond roughly to fields that are short and horizontally elongated or long and vertically elongated (green and red shading respectively). Examples of fields satisfying these criteria can be seen on the right (green area top row, red area bottom row). **b** Scatter graphs showing the relationship between field position in the mazes and field elongation. In both lattice mazes place field elongation increases further from the maze center (i.e. closer to the boundaries). Red lines represent the least squares line of best fit, red text outlines the result of a pairwise Pearson correlation on each data set. **c** Markers represent sessions, columns are as above. The relationship between place field elongation and time in the mazes. Text gives the result of independent pairwise Spearman's correlations. Different colors correspond to different rats. **d** Markers represent fields, the relationship between place field elongation and three cluster quality measures. Red text outlines the result of a pairwise Spearman correlation on each data set. Source data are provided as a Source Data file.

*Field orientation*

For more information on this analysis see Supplementary Methods: *Field orientation and size* and Fig. S8a. For each maze, the observed field orientation density map best correlates with the prediction made for that maze based on the maze axes (arena correlation with predicted arena, aligned and tilted maps: 0.74, 0.30 & -0.23 respectively, chance 99$^{th}$ percentile: 0.38; aligned correlation with predicted arena, aligned and tilted maps: 0.21, 0.92 & -0.32 respectively, chance 99$^{th}$ percentile: 0.41; tilted correlation with predicted arena, aligned and tilted maps: -0.07, -0.36 & 0.46 respectively, chance 99$^{th}$ percentile: 0.4; tilted (rotated) correlation with predicted arena, aligned and tilted maps: 0.17, 0.85 & -0.28 respectively, chance 99$^{th}$ percentile: 0.44). These effects can be seen in Fig. S8b.

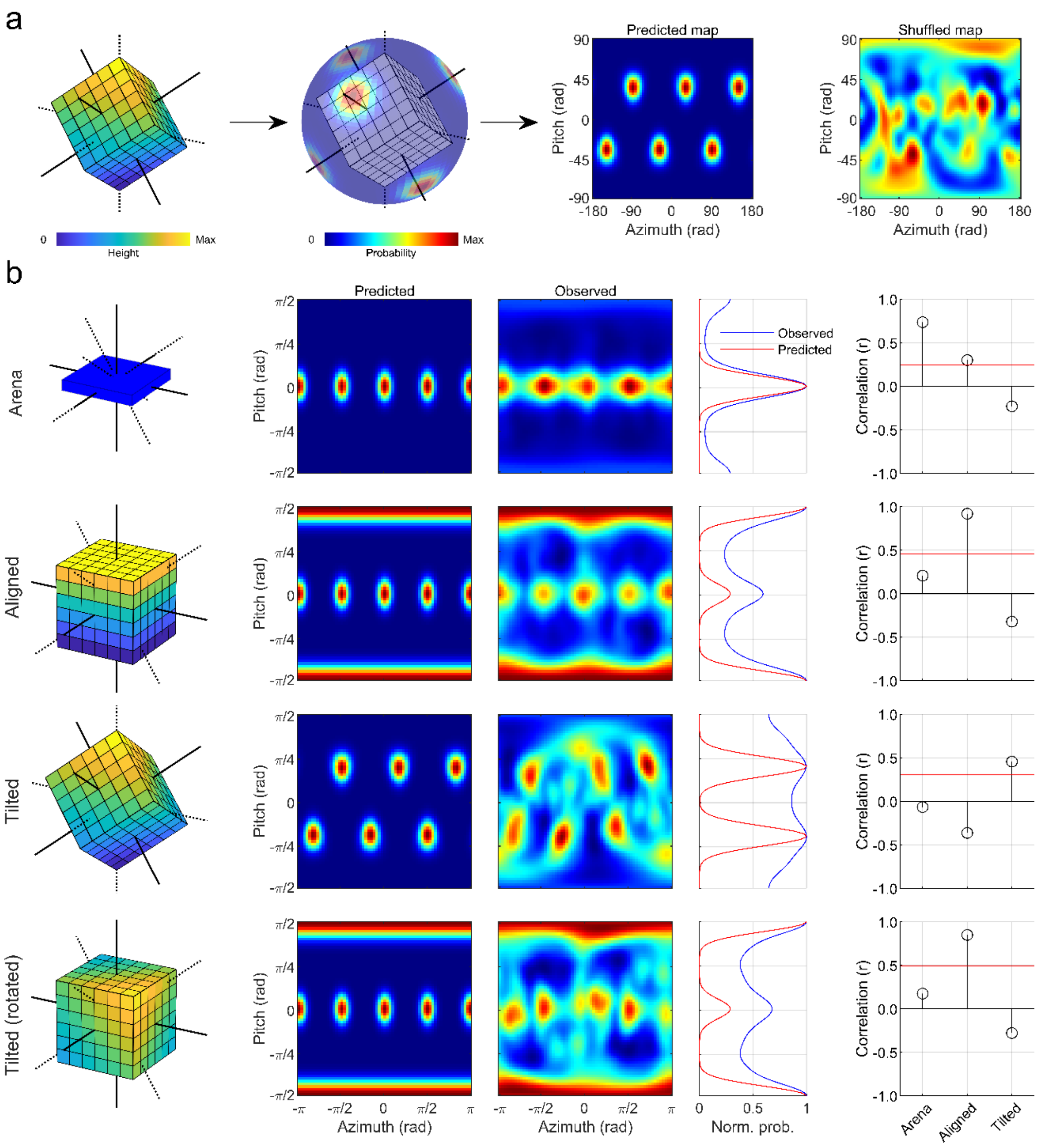


**Fig. S8:** Place field orientation in three dimensions. **a** Diagram demonstrating how place field orientations can be predicted from maze axes. The axes of the diagonal lattice (left) are projected onto a unit sphere and the Von-Mises Fisher density of these points are mapped (middle), this map can be projected cylindrically to form a 2D map (right). Shuffled maps can be generated in the same way from random spherical points (far right). **b** From top to bottom, rows show data for the open field arena, aligned lattice, tilted lattice and tilted lattice after rotating its data to mimic the aligned position. The first column shows a schematic of the maze. The second column shows the predicted probability of fields aligning with every possible three-dimensional orientation if they are parallel to the mazes axes (the walls, boundaries or lattice bars) in a 2D projection. The third column shows the observed pattern of results in each case, also as a 2D projection. The fourth column shows the summed, normalized probability of observing fields at each pitch angle in both the predicted and observed data. The last column shows the result of correlating the observed field density map with each possible prediction. Red lines show the upper $99^{th}$ percentile of a shuffle distribution (Supp. Methods: *Field orientation and size*).

*Autocorrelation, spatial information and binary morphology*

In the main text we report the result of an autocorrelation analysis where the central regions of the autocorrelation maps are extracted along each axis by interpolation. We found similar results when taking the median of central portions extracted from these autocorrelations. See Supp. Methods: *Autocorrelation and spatial information* for the distinction between these. Utilizing this approach we found that place cell firing rate maps in the aligned lattice were significantly more self-similar along the Z dimension (aligned X, Y & Z median autocorrelation: 0.05, 0.06 & 0.14; $\chi^2(2) = 115.5$, $p < .0001$, FT; X vs Y, $p = 0.97$, X vs Z and Y vs Z, $p < .0001$). In the tilted lattice there was a small but significant difference between the X and Y axes (tilted X, Y & Z median autocorrelation: 0.02, 0.04 & 0.04; $\chi^2(2) = 6.9$, $p = .032$, FT; X vs Y, $p = 0.026$, X vs Z and Y vs Z, $p > .50$) although none of the axes reached a correlation nearly as high as the Z axis in the aligned lattice ($\chi^2(3) = 271.6$, $p < .0001$, K-W; all comparisons to aligned Z, $p < .0001$). There were no differences between the A, B and C axes (tilted A, B & C median autocorrelation: 0.05, 0.06 & 0.06; $\chi^2(2) = 5.1$, $p = .079$, FT). These effects can be seen in Fig. S9c.

In addition we also employed a binary morphological approach, which utilizes thresholded ratemaps (Supp. Methods: *Binary morphology*). This method concentrates on the connectivity of neighboring voxels rather than firing rate or correlation. Through this approach we found that aligned lattice connectivity (see Fig. S18 for a graphical explanation) was near 33% for all three axes up to a distance of 7 voxels, meaning that voxels were equally connected to contiguous neighbors 7 voxels away along all three axes. However, after this point the connectivity begins to diverge significantly (main effect of axis: $F(2,9382) = 328.3$, $p < .0001$, $\eta_p^2 = 0.063$, interaction between axis and distance: $F(35,9382) = 17.6$, $p < .0001$, $\eta_p^2 = 0.061$, repeated measures ANOVA comparing effects of dimension and voxel distance on voxel proportion). Post-hoc tests confirm that each dimension differs from every other, with the Z-axis demonstrating higher connectivity at longer distances (X, Y & Z, mean proportion: 0.245, 0.297 & 0.457; all comparisons, $p < .0001$, pairwise comparisons with Bonferroni correction).

In the tilted lattice connectivity was also near 33% for all three axes at shorter distances but again this diverged significantly at longer distances (main effect of axis: $F(2,5766) = 17.1$, $p < .0001$, $\eta_p^2 = 0.006$, interaction between axis and distance: $F(36, 5766) = 2.2$, $p < .0001$, $\eta_p^2 = 0.014$, repeated measures ANOVA comparing effects of dimension and voxel distance on voxel proportion) although this effect is accompanied by much smaller effect sizes. Post-hoc tests confirmed that the A-axis demonstrated higher connectivity overall, while the B and C axes did not differ (A, B & C, mean proportion: 0.376, 0.310 & 0.314; A vs B and A vs C, $p < .0001$, B vs C, $p > .99$, pairwise comparisons with Bonferroni correction). However, in this maze all three axes remained much closer to equilibrium (33%) than in the aligned lattice. The same pattern of results was also obtained using raw values instead of proportions (data not shown). These effects can be seen in Fig. S9d.

Lastly, to investigate this reduced vertical resolution at the level of individual place fields we found the three orthogonal 1D Gaussians (parallel to the Cartesian axes) that best fitted each place field. In the aligned lattice these Gaussians differed in terms of their standard deviation (median X, Y & Z s.d.; 3.0, 2.7 & 4.1, $\chi^2(2) = 15.4$, $p = .0004$, $\eta_p^2 = .023$, K-W) with the vertical Gaussian (parallel to the Z-axis) best described by a larger standard deviation (X vs Z & Y vs Z, $p < .02$, X vs Y, $p > .92$). By contrast, fields in the tilted lattice could be adequately described by three Gaussians with equivalent standard deviation (median X, Y & Z s.d.; 4.2, 3.6 & 4.1, $\chi^2(2) = 5.0$, $p = .08$, $\eta_p^2 = .008$, K-W). However, the same analysis repeated with Gaussians parallel to the axes of the tilted lattice revealed a small but significant difference between the B and C axes, although this was accompanied by a small effect size (Median A, B & C s.d.; 3.5, 4.1 & 3.4, $\chi^2(2) = 7.5$, $p = .023$, $\eta_p^2 = .009$, K-W; A vs B & A vs C, $p > .23$, B vs C, $p = .02$). These effects can be seen in Fig. S9f.

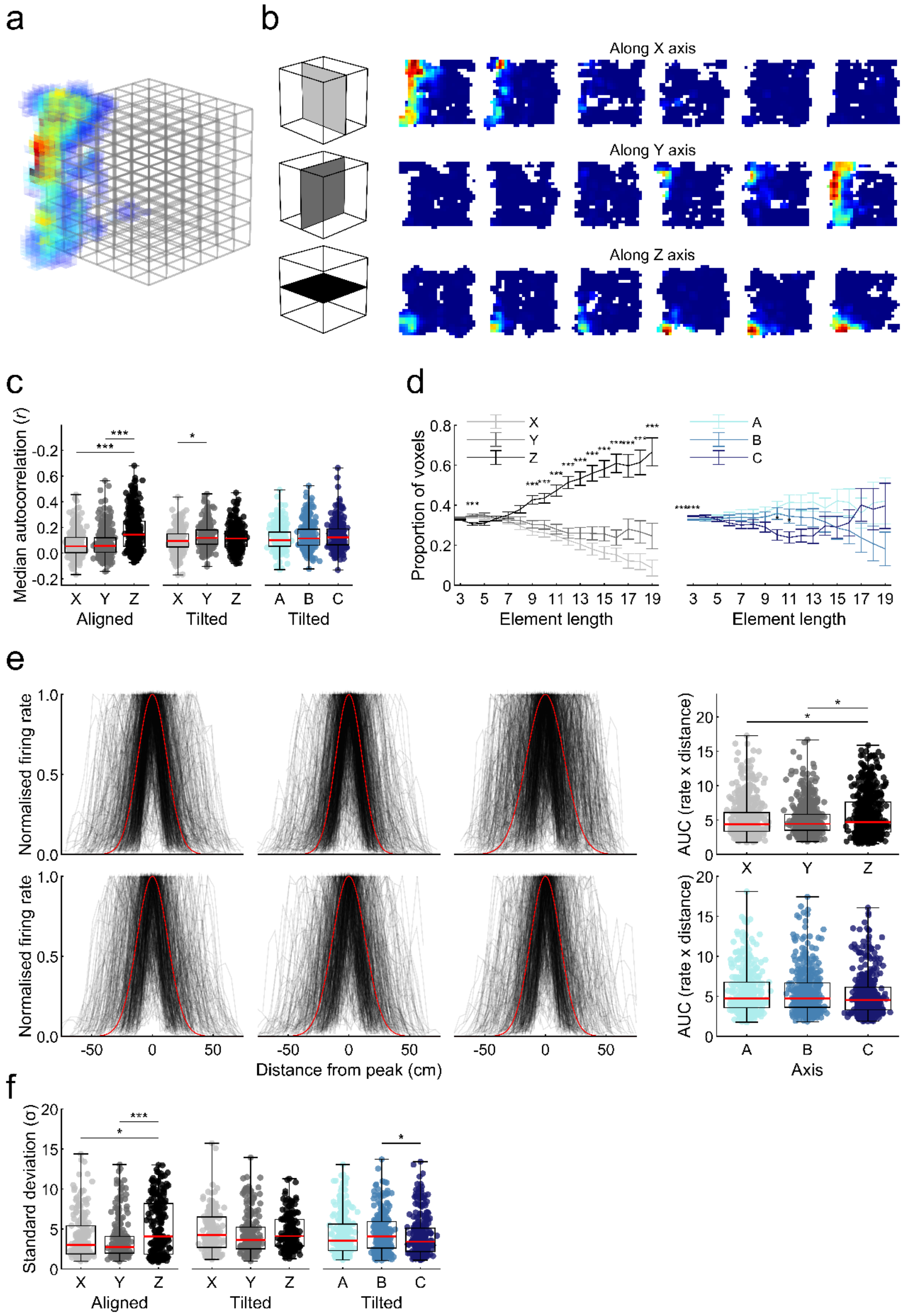
a
b
Along X axis
Along Y axis
Along Z axis
c
Median autocorrelation (r)
Aligned
Tilted
Tilted
d
Proportion of voxels
Element length
Element length
e
Normalised firing rate
Distance from peak (cm)
AUC (rate x distance)
Axis
f
Standard deviation (σ)
Aligned
Tilted
Tilted

**Fig. S9:** Spatial information in three dimensions. **a** An example ratemap of an elongated field recorded in the lattice maze. **b** The result of taking slices along each axis of the ratemap shown in **a**, so that each slice represents a layer of the lattice maze. **c** The median value in the central portion of each place cell's autocorrelogram (see Fig. S19 for a description). Values are shown for when the portion is taken parallel to the X, Y and Z-axes. Place cells generally have a higher median autocorrelation along the Z-axis, suggesting that lattice maze ratemaps are more self-similar along this axis. **d** The results of binary morphological analysis on thresholded firing rate maps for the aligned lattice (left) and tilted lattice (right). This analysis tests how many neighboring voxels each voxel is connected to (where connected means sharing one face). In the aligned lattice, voxels are more highly connected along the Z-axis, in the tilted lattice the maze axes are equivalent. **e** Black semi-transparent lines represent the peak-normalized firing rate distribution of every place field recorded in the aligned lattice (top row) and tilted lattice (bottom row) when projected onto the X,Y & Z axes (for the aligned lattice) or the A, B & C axes (for the tilted lattice). Red lines represent the mean Gaussian fitted to each population - at a width of 3 standard deviations. Note the slightly wider shape to the Z axis projections in the aligned lattice. The right box plots show the area under the curve (AUC, Matlab function *trapz*) of these distributions. Note the significantly higher values for Z in the aligned lattice. **f** The standard deviation of orthogonal 1-dimensional Gaussians fitted to place fields in each lattice maze. Source data are provided as a Source Data file.

*Comparison of firing properties between mazes*

See Supp. Methods: *Comparing activity between mazes* for an explanation of the analyses used here. The firing rate of place cells was significantly lower in the tilted lattice compared to the arena ($\chi^2(2)$ = 10.2, *p* = .0062, $\eta_p^2$ = .008, KW; arena vs aligned, *p* = .16, arena vs tilted, *p* = .0067, aligned vs tilted, *p* = .56, Fig. S10a). Spatial information content calculated on volumetric firing rate maps was also significantly lower in the tilted lattice than the other mazes ($\chi^2(2)$ = 11.8, *p* = .0027, $\eta_p^2$ = .009, KW; arena vs aligned, *p* > .99, arena vs tilted, *p* = .0058, aligned vs tilted, *p* = .0048, Fig. S10a), related to this sparsity was significantly higher in the tilted lattice than the other mazes ($\chi^2(2)$ = 11.0, *p* = .0041, $\eta_p^2$ = .008, KW; arena vs aligned, *p* > .99, arena vs tilted, *p* = .0072, aligned vs tilted, *p* = .0080, Fig. S10a). There was no relationship between the average elongation of fields in the arena and aligned lattice (*r* = 0.030, *p* = .64, Pearson’s pairwise correlation, Fig. S10b) but there was a weak negative correlation between the elongation of fields in the arena and tilted lattice (*r* = -0.177, *p* = .033, Pearson’s pairwise correlation, Fig. S10b). In the lattice mazes the lengths of place fields belonging to the same cell were not more similar than would be expected by chance (aligned vs chance: $\chi^2(1)$ = 0.9, *p* = .35, $\eta_p^2$ = .001, FT, tilted vs chance: $\chi^2(1)$ = 1.5, *p* = .22, $\eta_p^2$ = .003, FT, Fig. S10c) nor did fields belonging to the same cell share similar orientations at an above chance level (aligned vs chance: $\chi^2(1)$ = 0.0, *p* = .96, $\eta_p^2$ < .001, FT, tilted vs chance: $\chi^2(1)$ = 1.5, *p* = .22, $\eta_p^2$ = .003, FT, Fig. S10d). The number of fields expressed by cells was also not more consistent between the arena and lattice mazes than would be expected by chance (aligned vs chance: $\chi^2(1)$ = 0.0, *p* = .85, $\eta_p^2$ < .001, FT, tilted vs chance: $\chi^2(1)$ = 0.0, *p* > .99 $\eta_p^2$ < .001 FT, Fig. S10e).

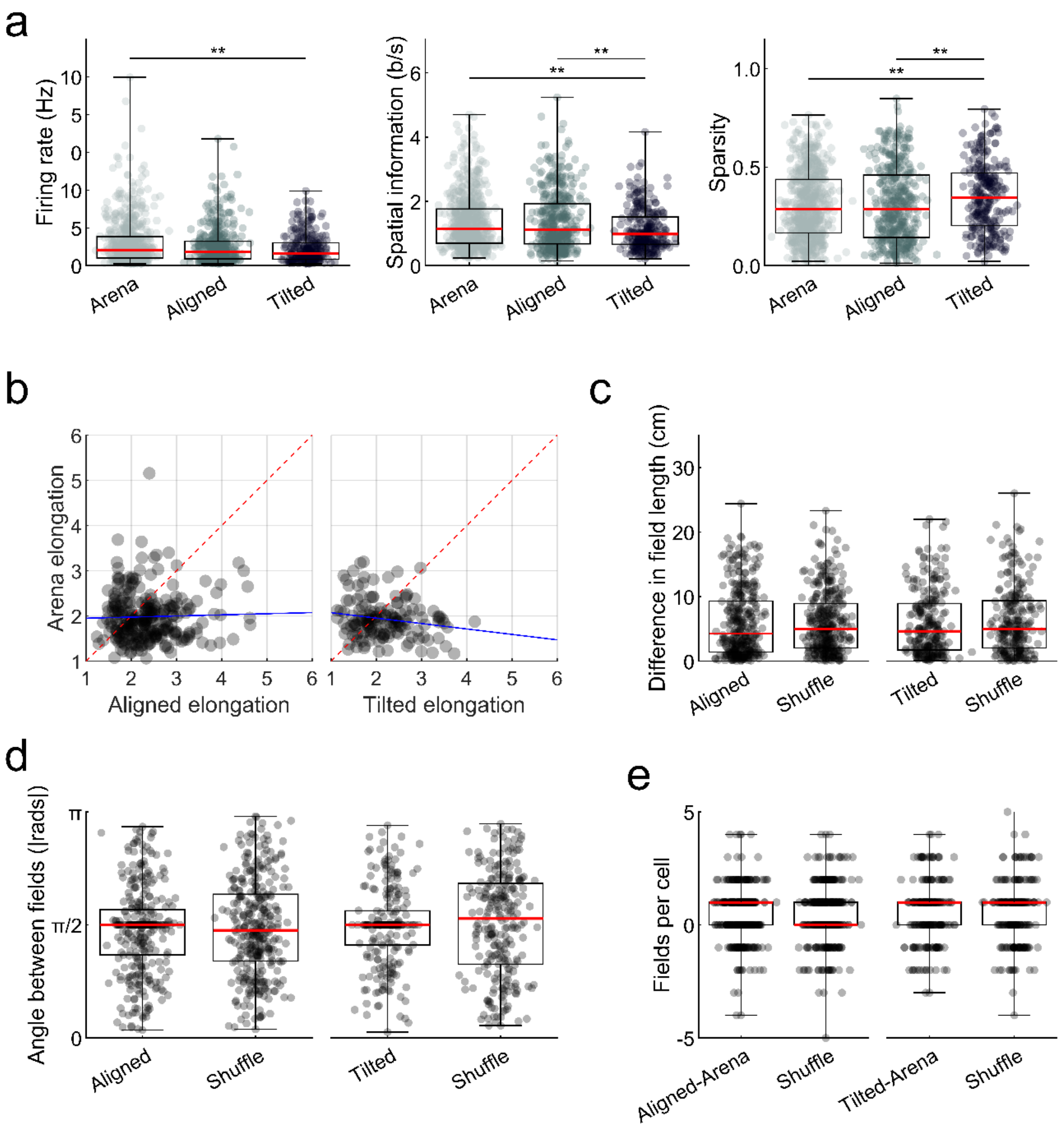


**Fig. S10:** Comparison of various firing and field characteristics between mazes. **a** Firing rate, spatial information content and sparsity of place cells recorded in the three mazes. **b** Relationship between field elongation in the arena and lattice mazes. Markers represent average values for cells, dotted red line denotes a linear correspondence, blue line shows the relationship resulting from a linear least squares fit. **c** Markers represent cells. For cells with multiple place fields in the aligned or tilted lattice, the average pairwise difference in major axis length of these fields. **d** Markers represent cells. For cells with multiple place fields in the aligned or tilted lattice, the average angle between their major axes. **e** Markers represent cells. For all place cells in the aligned or tilted lattice, the number of fields expressed in the lattice minus the number of fields expressed in the arena. Source data are provided as a Source Data file.

*Field stability*

We compared the first and second half of each maze session to determine the stability of place cell firing patterns (Fig. S11a, Supp. Methods: *Field stability*). These effects can be seen in Fig. S11b. Similar effects were observed when tilted lattice maze data were projected onto the Cartesian planes (XY projections: $z = 17.1$, $p < .001$, shuffle 95$^{th}$: 0.309, observed median: 0.274;  XZ projections: $z = 17.4$, $p < .001$, shuffle 95$^{th}$: 0.323, observed median: 0.275; YZ projections: $z = 17.0$, $p < .001$, shuffle 95$^{th}$: 0.279, observed median: 0.259; whole volume: $z = 20.6$, $p < .001$, shuffle 95$^{th}$: 0.125, observed median: 0.146, statistics refer to right-tailed Wilcoxon rank sum tests).

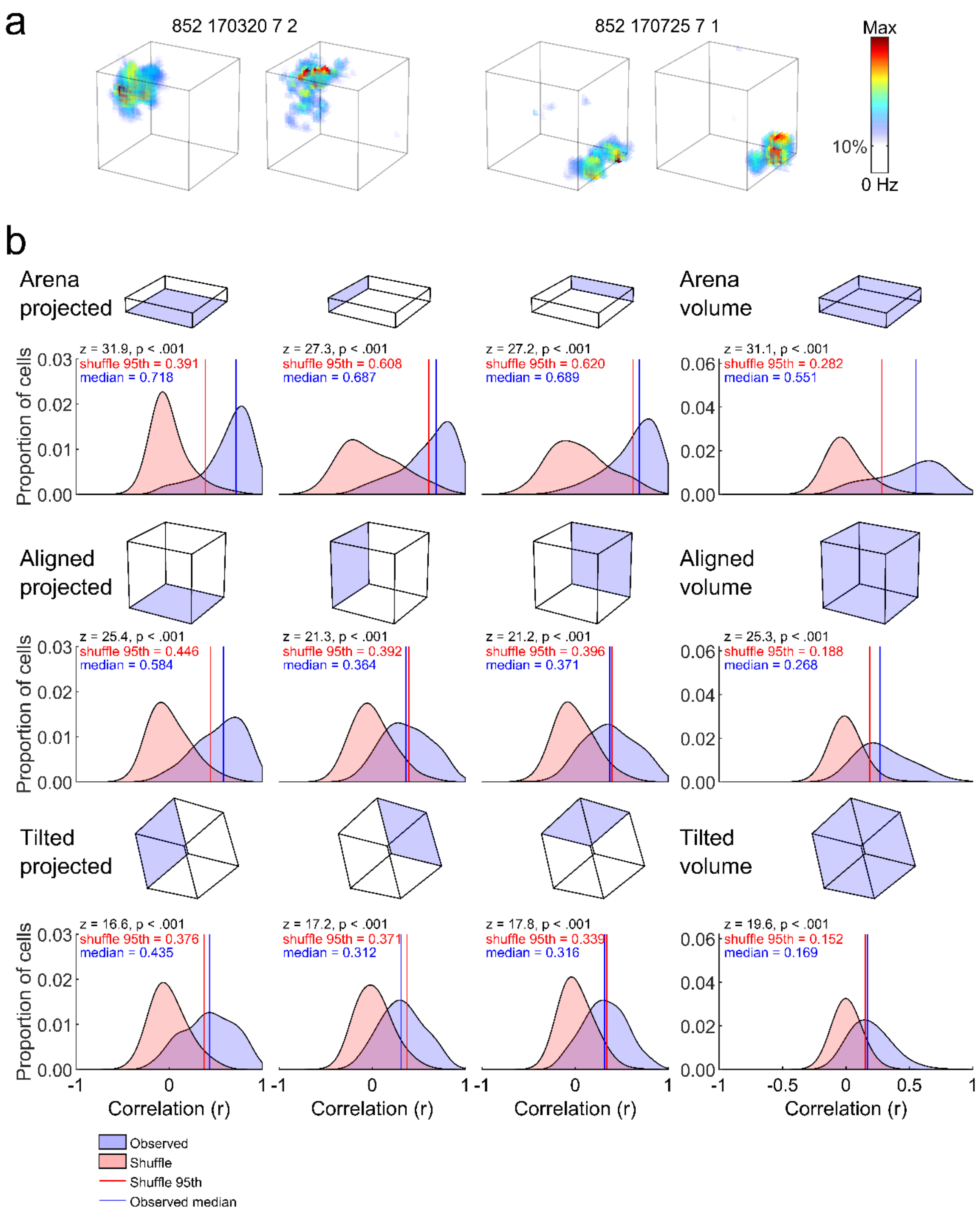
a
852 170320 7 2
852 170725 7 1
Max
10%
0 Hz
b
Arena projected
Arena volume
Aligned projected
Aligned volume
Tilted projected
Tilted volume
Proportion of cells
Correlation (r)
z = 31.9, p < .001
shuffle 95th = 0.391
median = 0.718
z = 27.3, p < .001
shuffle 95th = 0.608
median = 0.687
z = 27.2, p < .001
shuffle 95th = 0.620
median = 0.689
z = 31.1, p < .001
shuffle 95th = 0.282
median = 0.551
z = 25.4, p < .001
shuffle 95th = 0.446
median = 0.584
z = 21.3, p < .001
shuffle 95th = 0.392
median = 0.364
z = 21.2, p < .001
shuffle 95th = 0.396
median = 0.371
z = 25.3, p < .001
shuffle 95th = 0.188
median = 0.268
z = 16.6, p < .001
shuffle 95th = 0.376
median = 0.435
z = 17.2, p < .001
shuffle 95th = 0.371
median = 0.312
z = 17.8, p < .001
shuffle 95th = 0.339
median = 0.316
z = 19.6, p < .001
shuffle 95th = 0.152
median = 0.169
Observed
Shuffle
Shuffle 95th
Observed median

359

**Fig. S11:** Stability of recording between first and second halves of each maze recording. **a** Ratemaps for two example cells from the first (left map) and second half (right map) of an aligned lattice recording. **b** Distributions of observed correlation scores between the first and second halves (blue shaded area) and between 1000 shuffled first and second halves (red shaded area). Black text gives the result of a right-tailed Wilcoxon rank sum test comparing the distributions. Red lines denote the 95th percentile rank position in the shuffled distribution, blue lines denote the median position in the observed distribution. The top row shows the result of this analysis when comparing the 1st and 2nd half of arena recordings after projecting all data onto the XY, XZ and YZ planes respectively; and when volumetric firing rate maps are compared without projection. The second row shows the same for the aligned lattice. The third row shows the same for the tilted lattice but here projections are made along the AB, AC and BC planes instead; values for the Cartesian planes are given in text. Source data are provided as a Source Data file.

*Theta-speed relationships differed between the environments*

Theta power was significantly higher in the aligned configuration (Fig. S12a&c; arena, aligned & tilted median power: 264, 421 & 252 $\mu V^2/Hz$; $\chi^2(2) = 9.9$, *p* = .007, $\eta_p^2$ = 0.09, KW; arena vs aligned: *p* = .015, arena vs tilted: *p* > .99, aligned vs tilted: *p* = .036), and increased with running speed in the arena but not the lattice (Fig. S12d; speed/power correlations: arena, aligned & tilted median r: 0.94, 0.69 & 0.76; $\chi^2(2) = 64.4$, *p* < .0001, $\eta_p^2$ = 0.57, KW; arena vs aligned: *p* < .0001, arena vs tilted: *p* < .0001, aligned vs tilted: *p* = .95). Fitted *b* parameters (Supp. Methods: *Running speed analyses*) were significantly higher for the arena than the lattice mazes (arena, aligned & tilted median *b*: 0.34, 0.01 & 0.01; $\chi^2(2)$ = 62.8, *p* < .0001, $\eta_p^2$ = 0.55, KW; arena vs aligned: *p* < .0001, arena vs tilted: *p* < .0001, aligned vs tilted: *p* > .99) indicating a linearly increasing relationship in the open field data but a downwardly curved or approximately flat one in the lattice data.

Theta frequency was significantly lower in the tilted configuration (Fig. S12b-c, arena, aligned & tilted median frequency: 8.99, 9.21 & 8.61 Hz; $\chi^2(2)$ = 19.5, *p* < .0001, $\eta_p^2$ = 0.55, KW; arena vs aligned: *p* = .055, arena vs tilted: *p* = .015, aligned vs tilted: *p* < .0001) but increased with running speed equally in all environments (Fig. S12e; arena, aligned & tilted median r: 0.93, 0.91 & 0.91; $\chi^2(2)$ = 2.7, *p* = .26, $\eta_p^2$ = 0.02, KW). Fitted *b* parameters were close to zero in all cases, with the aligned lattice values significantly lower than the other two mazes (arena, aligned & tilted median *b*: 0.01, 0.01 & 0.01; $\chi^2(2)$ = 12.5, *p* = .002, $\eta_p^2$ = 0.11, KW; arena vs aligned: *p* > .99, arena vs tilted: *p* = .0052, aligned vs tilted: *p* = .002) indicating approximately flat relationships in all cases.

When comparing the three mazes firing rates were found to be modulated by running speed (F(24,23072) = 12.8, *p* < .0001, $\eta_p^2$ = 0.013) but this differed significantly between environments (F(2,23072) = 1147.0, *p* < .0001, $\eta_p^2$ = 0.09) with a significant interaction between speed and environment (F(48,23072) = 2.3, *p* < .0001, $\eta_p^2$ = 0.005). Post-hoc tests confirmed that every maze differed significantly from every other maze (*p* < .0001 in all cases, Fig. S13a-b).

Firing rates were generally lower in the lattice mazes but only the open field and tilted lattice differed significantly (arena, aligned & tilted median firing rates: 0.83, 0.73 & 0.64 Hz; $\chi^2(2) = 10.2$, $p < .0062$, $\eta_p^2 = 0.008$, KW; arena vs aligned: $p = .16$, arena vs tilted: $p = .0067$, aligned vs tilted: $p = .56$, Fig. S13c). These reduced firing rates did not negatively affect the accuracy of the speed-rate linear regressions as fit error was significantly smaller in the lattice maze environments (arena, aligned & tilted sum of squared errors: 1.22, 0.74 & 0.67; $\chi^2(2) = 207.7$, $p < .0001$, $\eta_p^2 = 0.18$, KW; arena vs aligned: $p < .0001$, arena vs tilted: $p < .0001$, aligned vs tilted: $p = .042$). Dwell time was significantly inversely related to running speed ($F(24,1225) = 126.0$, $p < .0001$, $\eta_p^2 = 0.71$) but this did not differ significantly between environments ($F(2,1225) = 0.2$, $p = .82$, $\eta_p^2 < 0.001$) nor was there a significant interaction between the two ($F(48,1225) = 1.2$, $p = .17$, $\eta_p^2 = 0.045$, Univariate ANOVA comparing effects of speed and environment on dwell time, Fig. S13d).

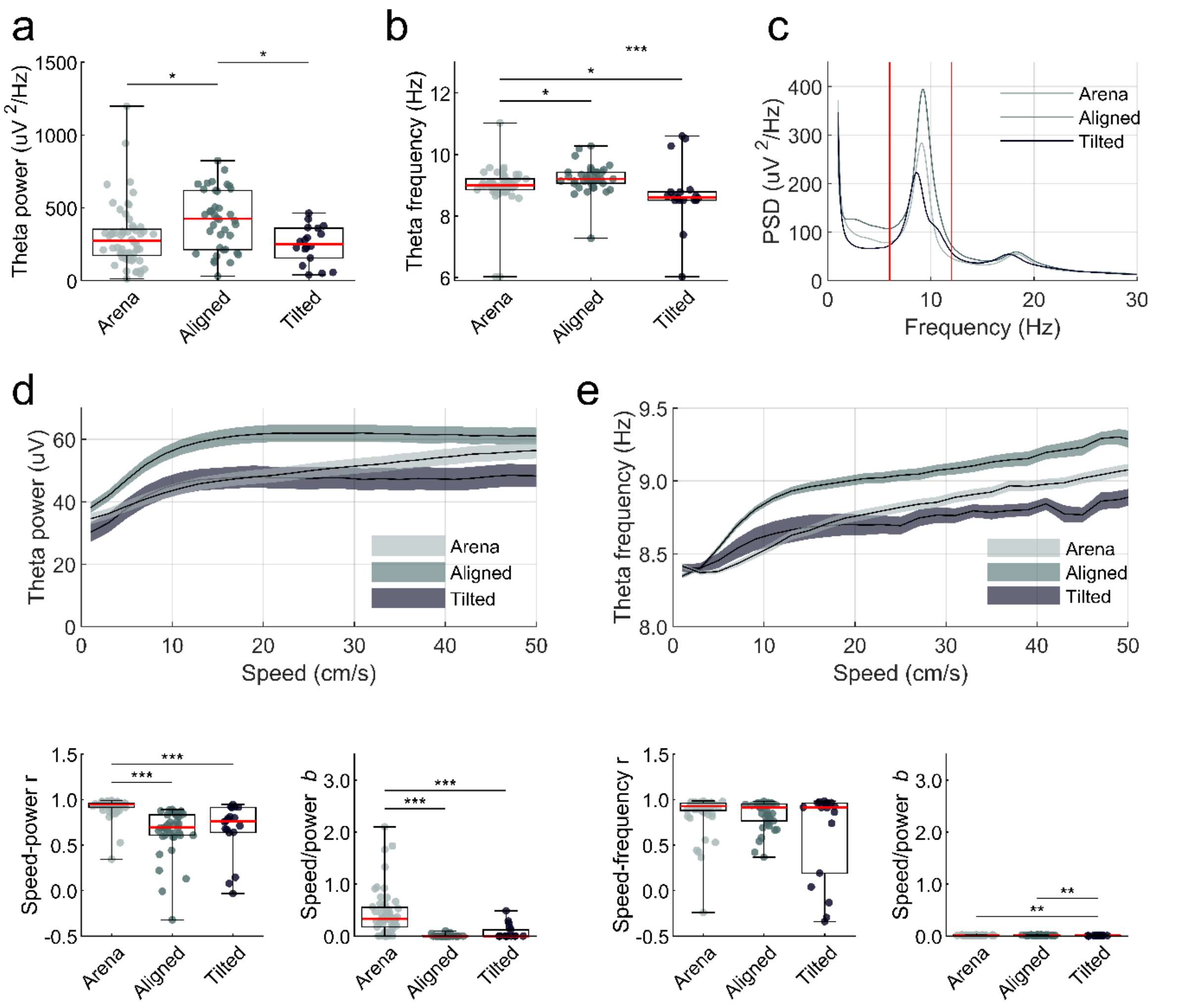


**Fig. S12:** The relationship between theta and running speed. **a** Markers represent sessions. The maximum power found in the theta band (6-12Hz) of the Welch power spectral density estimate (PSD) calculated using whole session LFPs. **b** Markers represent sessions. The frequencies associated with the maximum power in the theta band. **c** The mean PSD in each maze, averaged across sessions. Red lines denote the frequency band associated with theta rhythm. **d** For each maze, the mean and SEM theta power (amplitude of the Hilbert transform) observed at various running speeds, averaged across sessions. Below this are boxplots where markers represent sessions. These show the Pearson's correlation between power and speed (left) and the *b* parameter extracted from each session's speed-power curve (right). **e** Same as **d** but for frequency (derivative of the phase of the Hilbert transform). Source data are provided as a Source Data file.

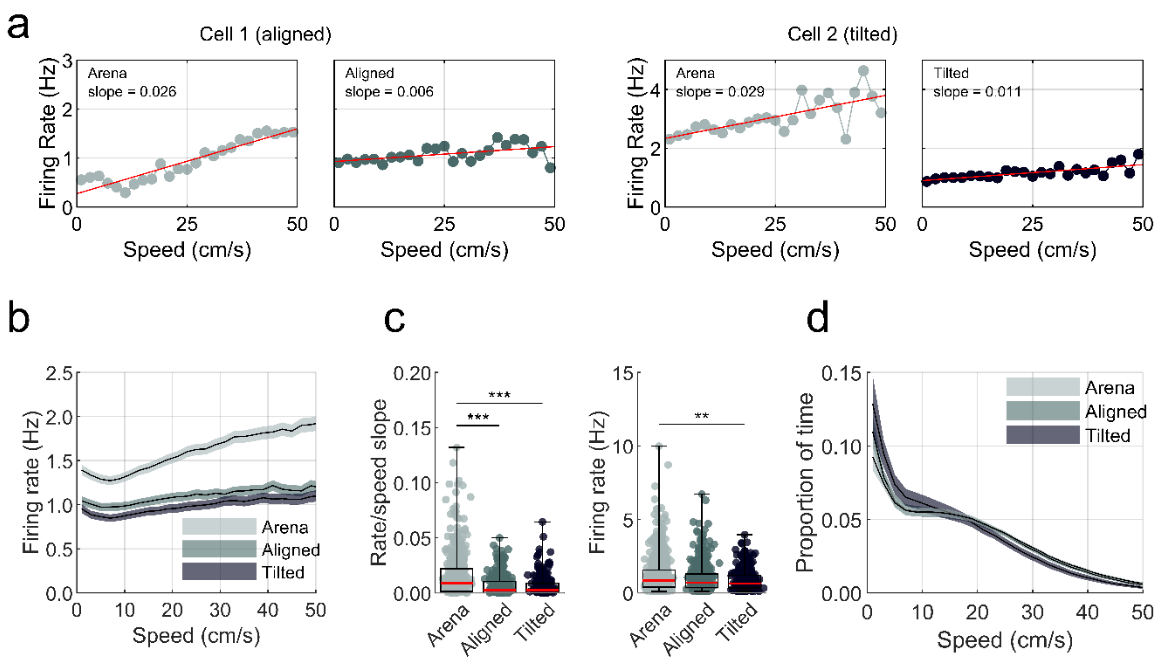


**Fig. S13:** The relationship between place cell activity and running speed. Similar relationships were also observed in non-spatially modulated pyramidal cells and interneurons (data not shown). **a** Firing rate by speed profiles for two place cells. Left two plots are for a cell recorded in the open field and aligned lattice respectively. Right two plots are for a cell recorded in the open field and tilted lattice respectively. Markers show the cell's average firing rate at that running speed, calculated across the whole session, red lines show the result of a linear least squares line of best fit. We extract two main features from these fits, the first is the y-axis intercept and the second is the slope of the line. Both cells show a reduced slope and thus speed modulation in the lattice mazes. **b** The mean and SEM firing rate of cells at various running speeds, averaged across place cells. **c** Markers represent place cells. Left) for each cell we extracted the slope of the line of best fit (red lines in **a**). These slopes are significantly lower in the lattice mazes when compared to the open field. Right) The overall mean firing rate of cells in the three mazes. **d** The mean and SEM proportion of time animals spent moving at various running speeds, averaged across sessions. Source data are provided as a Source Data file.

*The relationship between theta and place cell spiking differed between the environments*

Theta modulation of individual cells, assessed using spike autocorrelations, was significantly lower for place cells in the tilted lattice (arena, aligned & tilted median theta index: 0.26, 0.28 & 0.21 Hz; $\chi^2(2) = 38.4$, $p < .0001$, $\eta_p^2 = 0.033$, KW; arena vs aligned: $p > .99$, arena vs tilted: $p < .0001$, aligned vs tilted: $p < .0001$, Fig. S14a). Place cells usually burst at a rate slightly faster than theta due to precession of their firing phase [1] which possibly reflects odometry [2]. Overall intrinsic frequency of cells in the open field was faster than the global theta rhythm (median intrinsic vs. global frequency: 9.12 vs. 8.92 Hz, z = -10.7, $p < .001$, WSR; 275 cells or 79% faster than global, $\chi^2(1) = 115.8$, $p < .0001$, Chi-square test of expected proportions). The same was also true in the aligned lattice (intrinsic vs. global frequency: 9.21 & 9.07 Hz, z = -2.4, $p = .016$, WSR; 208 cells or 58% faster than global, $\chi^2(1) = 9.4$, $p = .002$, Chi-square test of expected proportions) but this proportion was significantly lower than in the open field ($\chi^2(1) = 36.0$, $p < .0001$, Chi-square test of expected proportions). Lastly, in the tilted lattice, overall burst frequency was faster than theta (median intrinsic vs. global frequency: 8.82 vs. 8.51 Hz, z = -6.8, $p < .001$, WSR; 170 cells or 78% faster than global, $\chi^2(1) = 66.9$, $p < .0001$, Chi-square test of expected proportions) and this proportion was more similar to the open field ($\chi^2(1) = 0.1$, $p = .78$, Chi-square test of expected proportions) but this was founded on a much smaller number of sessions (arena, aligned & tilted total sessions: 50, 34 & 16). These effects can be seen in Fig. S14b.

Place cells were similarly modulated by theta phase in the open field arena and aligned lattice but less strongly modulated in the tilted lattice (arena, aligned & tilted median Rayleigh vector lengths: 0.14, 0.11 & 0.07; $\chi^2(2) = 47.1$, $p < .0001$, $\eta_p^2 = 0.01$, KW; arena vs aligned: $p = .62$, arena vs tilted: $p < .0001$, aligned vs tilted: $p < .0001$). Nevertheless, in all three mazes place cells exhibited a preference for a specific phase of theta at the population level (arena, aligned & tilted Rayleigh vector lengths: 0.53, 0.54 & 0.50, $p < .0001$ in all cases, Fig. S14c).

However, these distributions differed significantly (arena vs aligned: k = 77692, $p$ = .003; arena vs tilted: k = 47963, $p$ = .003, aligned vs tilted: k = 51130, $p$ = .003, two-sample Kuiper tests with Bonferroni correction) and cross-correlation confirms that the aligned lattice preferred phase was significantly earlier in phase than the arena (arena & aligned circular mean: -1.84 & -2.29 rad, correlation lag = -0.4 rad) while the tilted lattice was significantly later in phase than both of these (arena & tilted circular mean: -1.84 & -0.67 rad, correlation lag = +1.2 rad; aligned vs tilted correlation lag = +2.0 rad). These effects can be seen in Fig. S14c.

476

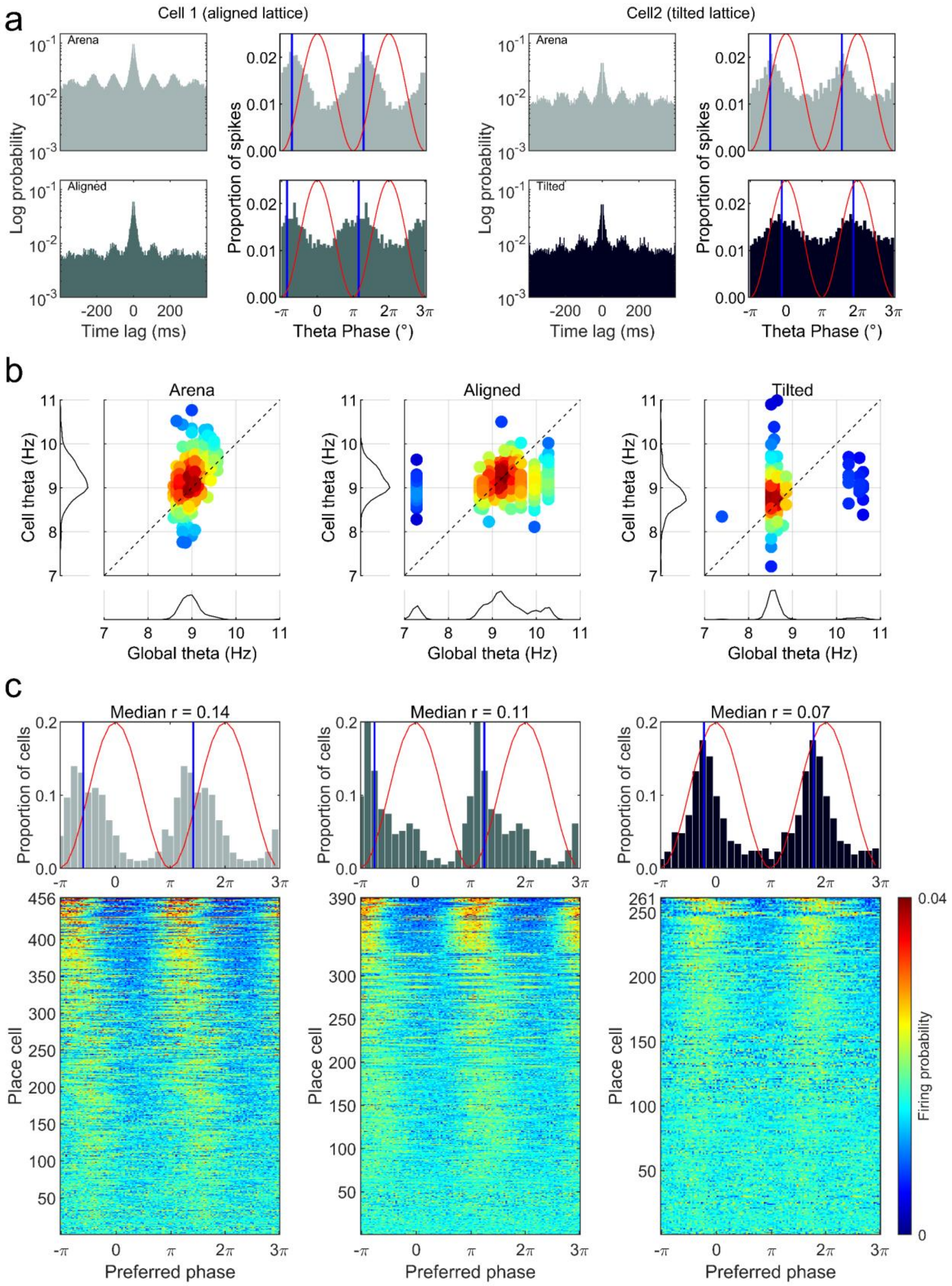

a
Cell 1 (aligned lattice)
Cell2 (tilted lattice)
Arena
Aligned
Tilted
Log probability
Proportion of spikes
Time lag (ms)
Theta Phase (°)
b
Arena
Aligned
Tilted
Cell theta (Hz)
Global theta (Hz)
c
Median r = 0.14
Median r = 0.11
Median r = 0.07
Proportion of cells
Place cell
Preferred phase
Firing probability

**Fig. S14:** Place cell activity and the theta oscillation. **a** Spike probability properties of two place cells. Left four plots are for a cell recorded in the open field (top row) and aligned lattice (bottom row). Right four plots are for a cell recorded in the open field (top row) and tilted lattice (bottom row). For each maze the left plot shows the 400ms spike autocorrelogram (1ms bins) and the right plot shows the spike-theta phase histogram. Red lines trace the amplitude of a scaled theta wave at every phase, blue lines show the circular mean of the phase angles. **b** Markers represent cells, color denotes marker density. Scatter plots showing the frequency of intrinsic theta modulation (extracted from the spike autocorrelograms) vs the overall frequency of global theta (extracted from the power spectral density estimate of the LFP). Markers above the dashed line represent cells firing at a rate faster than global theta, an indicator of phase precession. **c** Histograms showing the preferred theta phase (location of circular mean or blue lines in **a**) of all place cells. Red lines trace the amplitude of a scaled theta wave at every phase, blue lines show the circular mean of the preferred phases. Shown below each plot is the firing probability of all place cells relative to theta phase. Each row represents a cell and rows are sorted from top to bottom by the Rayleigh vector length of the cell's phase angles (i.e. the strength of phase-locking) from strong to weak. Source data are provided as a Source Data file.

*Histology*

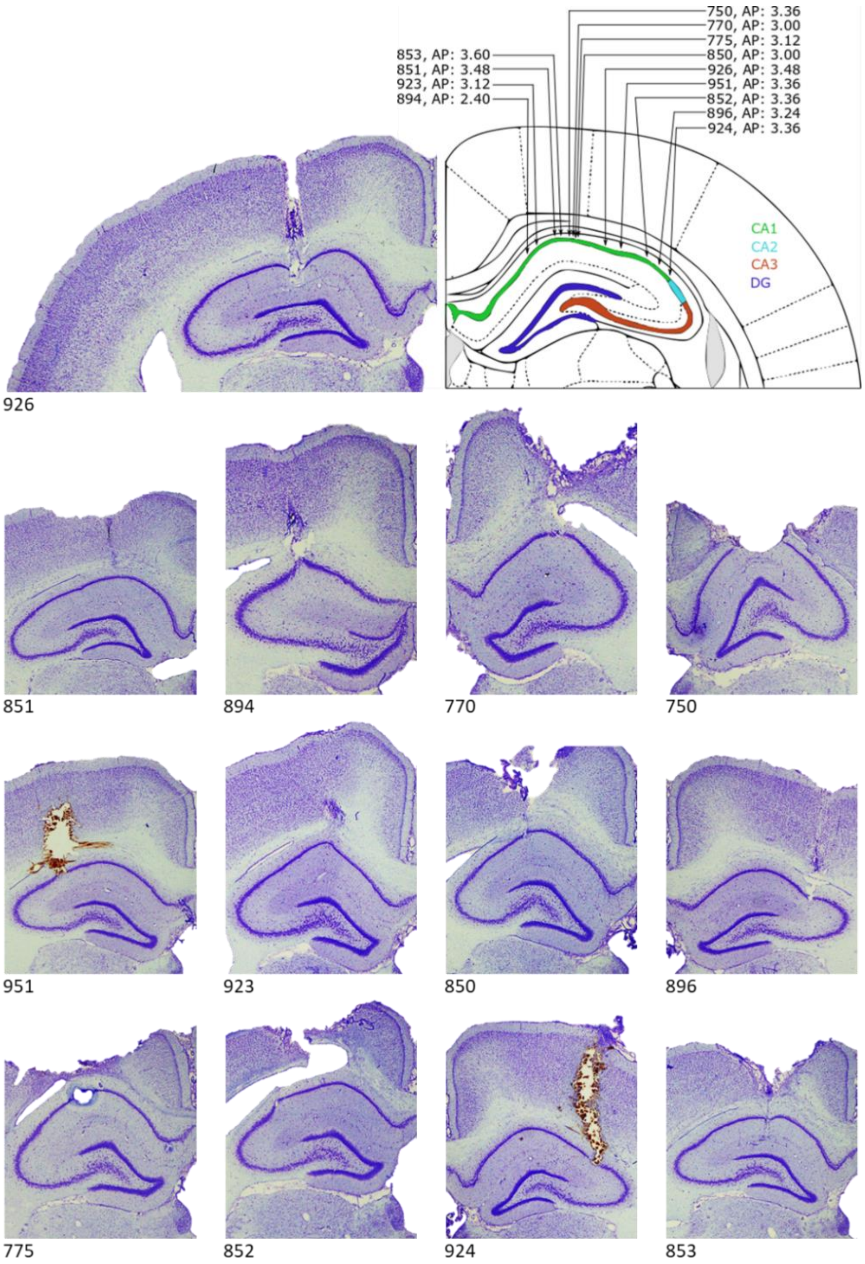


**Fig. S15:** Histological sections showing electrode tracks. Following fixation and sectioning, brains were imaged and recording electrode paths were reconstructed to determine the locations of recorded cells. In all animals the electrode tracks descended into dorsal CA1 and crossed the pyramidal cell layer without reaching dentate gyrus (DG) or CA3 below. Top: example histological slice (left) and the histological map corresponding to our intended coordinates (AP: -3.48, ML: ±2.4 & DV: -1.5mm). Arrows show the estimated trajectory of each animals' electrode bundle, annotations give the animal number and the estimated AP of the electrode track. Bottom: smaller hippocampal sections for every animal which best show the electrode bundle track. The proportion of data obtained from each animal can be seen in Table S2. Animal 775 received an electrolytic lesion under anesthesia prior to perfusion which can be seen as a small hole and surrounding tissue discoloration. Animals 951 and 924 lost their drive prior to perfusion resulting in staining of the electrode tracks. Source data are provided as a Source Data file.

### Supplementary methods

*Apparatus*

All experiments were conducted in the same room (3.2×2.1×2.2m) under moderately dimmed light conditions. Three of the room walls were covered with black material to aid position tracking; on two of these walls were large high-contrast cues (1.5×1.2m cardboard sheet and a 1×1.7m yellow plastic sheet). The last wall was covered with white material (2.2×2.2m white cotton). The floor of the room was covered with black anti-static linoleum flooring. We used three pieces of experimental apparatus; the first was a square open field environment ('arena'), the second was a cubic lattice composed of horizontal and vertical climbing bars ('aligned' lattice), the third was the same lattice rotated 45° around its Z-axis and 54.74° around its X-axis so that two of its vertices were vertically aligned ('tilted' lattice).

The arena was a 1.8×1.8m square high-walled wooden enclosure, composed of four 1.8×0.65m matte black painted walls. This enclosure was placed directly on the black linoleum flooring of the room. The top edges of these walls were covered with large corrugated tubing to prevent the rats from exploring this area. The bottom edge of this square was highlighted with a strip of 50% grey paint. One 0.45×0.65m matte white wooden cue was affixed to one wall of this enclosure; the position of this cue remained the same throughout the experiment. Rats were recorded freely foraging in the arena for randomly dispersed flavored puffed rice (CocoPops, Kelloggs, Warrington, UK).

The cubic lattice maze (Fig. 1) was constructed from a children's toy-set (Quadro, Hamburg, Germany). Hollow cubes were created by attaching red plastic tubes (length: 150mm, diameter: 10mm) using 6- or 4-way connectors (each 10mm wide). These cubes were then assembled into a 6×6×6 cubic maze (0.97×0.97×0.97m). The maze was raised 0.45m above the ground, initially on black metal stools but later on a narrow wooden frame. To encourage exploration, malt paste (GimCat Malt-Soft Paste, H. von Gimborn GmbH) was affixed to bars of the lattice by the experimenter. This paste was spread evenly throughout the maze, midway along bars, equally between horizontal and vertical bars and reapplied every 15 minutes. This maze could be placed on one side, with the bars running vertically and horizontally; we refer to this as the 'aligned' configuration. Alternatively, the maze could be rotated (45° around its Z-axis and 54.74° around its X-axis) so that two vertices were vertically aligned – essentially standing the lattice vertically on one corner. We refer to this as the 'tilted' lattice configuration (Fig. 1).

*Recording setup and procedure*

Single unit activity was observed and recorded using a custom built 64-channel recording system (Axona, St. Albans, UK). Mill-Max connectors built into the rat's microdrive were attached to a wireless headstage (custom 64-channel, W-series, Triangle Biosystems Int., Durham, NC). Analog signals were transmitted to a wireless base station via dual receiver antennae situated approximately 1m above the maze environments. Unfiltered signals were sampled at 50 kHz, amplified 100 times and transmitted at approximately 3.375 GHz (300 µW at 3m). They were then passed to an Axona pre-amplifier where they were amplified a further 100 times. The signal was then passed to a system unit and for single unit recording the signal was band-pass (Butterworth) filtered between 300 and 7000 Hz. Signals were digitized at 48 kHz and could be further amplified 10–40 times at the experimenter's discretion. For LFP recording a 4.8 kHz signal was saved as above which was then band-pass filtered between 6 and 12 Hz ($4^{th}$ order butterworth filter, Matlab *butter* and *filtfilt*) for theta analyses described below. The position of the animal was recorded using four wide-angle infrared LEDs (Osram Opto SFH 487P, 880nm) fixed to and powered by the wireless headstage. Five infrared sensitive CCTV cameras (Samsung SCB-5000P) tracked the animal's position at all times (Supp. Methods: *Trajectory reconstruction*). Tracking artefacts and reflections were not observed during recording or in the trajectory data (Fig. S17 and Supplementary Video S4).

After recovery from surgery, rats were screened for single unit activity and for the presence of theta oscillations once or twice a day, five days a week. Screening was performed in the open field apparatus, after which rats were given approximately equivalent experience freely foraging on the lattice maze. Once the presence of place cells was confirmed rats were recorded in the experimental apparatus.

In these sessions, rats were recorded for a minimum of 18 minutes in the open field environment and until they had sufficiently explored the environment (median, min & max session time: 20.1, 18.0 & 30.7 minutes). Without unplugging the wireless headstage, they were then removed and allowed to rest in an opaque, lidded box for approximately 10 minutes with access to drinking water. During this time, the open field environment was dismantled and replaced with the lattice maze in one of the two configurations described above. Rats were then placed on the bottom layer of the lattice maze (or the bottom front face of the diagonal lattice) and explored this environment for a minimum of 45 minutes and until they had sufficiently explored the environment (median, min & max session time: 60.2, 41.1 & 94.2 minutes). When this was complete, the rats were returned to the opaque box as before and the open field was restored. For a subset of recordings (41 or 71.9% of sessions) rats were then recorded for a further minimum of 16 minutes in the open field and until they had sufficiently explored the environment (median, min & max session time: 20.1, 10.9 & 30.9 minutes). During recordings the experimenters monitored progress from a connected room which housed the recording equipment and computers and was separated from the experimental room by a black opaque curtain.

At the end of the recording session, the animals were removed from the apparatus and the electrodes were lowered by at least 20 μm in order to maximize the chance of recording from a different population of cells on the following day. No attempt was made to track cells across days, and thus a subset of the cells may have been recorded on more than one session, although inspection of the cluster space did not suggest that this was the case and almost identical results to those reported in main text were also observed when only analyzing one session per animal (the session with the most place cells). These analyses can be replicated using the provided data set and code. Rats were tested until cells were no longer observed (median, min & max: 3, 1 & 14 sessions).

*Trajectory reconstruction*

Each animal's movements were monitored using five infrared CCTV cameras (Samsung SCB-5000P) mounted at the four corners of the room, with one camera directly above the environment. These tracked the position of four light-emitting diodes connected to the head-stage on the rat's head. Its position was then tracked, in real time, using custom software (DacqTrack, Axona, St. Albans, UK) at a 25Hz sampling frequency. The data from these cameras was synchronized with neural data using a pulsed optic interface – each camera monitored a 1Hz TTL initiated light source, controlled by the recording system, which allowed accurate, offline synchronization. The onset of these light pulses was used to continually re-align the position data using nearest neighbor interpolation (Matlab function *interp1*).

The rat's 3D position was then reconstructed using the direct linear transform algorithm [3], applied to the data from all five cameras, in pairs. Briefly, these cameras were first calibrated in order to reverse any distortion introduced by their optical elements (Matlab functions *estimateCameraParameters, undistortImage and undistortPoints*). We then imaged the same checkerboard pattern with each camera and used its 3D pose to calculate the distance and orientation of each camera relative to it and thus to each other (Matlab functions *extrinsics and cameraMatrix*). Using this information, we constructed a fundamental matrix. If $x$ are some points viewed by camera 1 and $x'$ are the same points viewed by camera 2, the fundamental matrix, $F$, represents the relationship between points $x$ and $x'$:

$$x_i' F x_i = 0$$

This relationship can be used to triangulate any given pair of points imaged by two cameras into three-dimensional space [3]. For each recording session we reconstructed the animal's path using every possible pair of cameras (Matlab function *triangulate*) and we then combined these reconstructions into one single trajectory. This was achieved by taking the weighted mean of each point, where the weighting was the reliability of the point's estimated location. Reliability was assessed using each point's reprojection error; after triangulation each point was projected back into both camera images; the reprojection error was then calculated as the distance between the original and reprojected position of the point.

In this setup, the rat need only be viewed by two cameras at any one time for a successful reconstruction, allowing for near continuous tracking even in cluttered, complex environments such as the lattice maze. For the lattice sessions reported here, rats were in view of at least two cameras for an average of 98.0% (geometric mean, geometric SD = 3.3%) of the session, this value was similar for the open field sessions (geometric mean = 97.6%, geometric SD = 2.5%). Based on reprojection errors, we estimate a median error in reconstruction of just 1.8 mm (median absolute deviation = 1.1mm). Camera trajectory reconstructions from different pairs of cameras were within a median of 13.8mm agreement (median absolute deviation = 7.4mm) of the weighted mean trajectory. Our cameras were extremely stable; however, re-calibrations were conducted once every two to four weeks to ensure continued reconstruction accuracy. For segments of missing tracking data, we simultaneously interpolated and smoothed the existing data using an unsupervised, robust, discretized, n-dimensional spline smoothing algorithm (Matlab function *smoothn* [4,5]). We did not observe reflections from the maze surfaces being tracked during recording, indeed the recording backdrop and lighting were specifically designed to eliminate tracking artefacts such as this. Examples of reflection-less tracking can be seen in Fig. S16 and Supplementary Video S4.

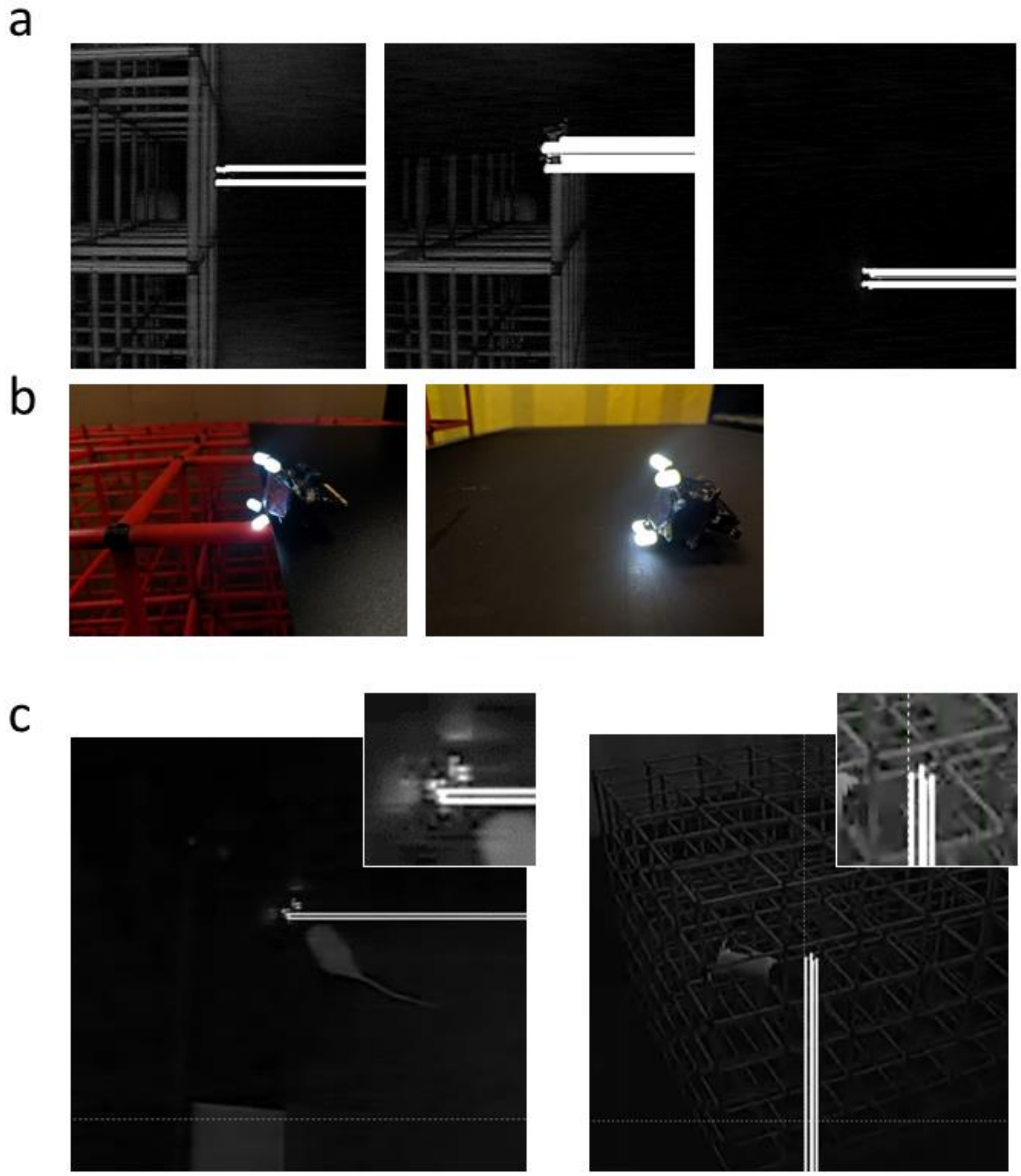


**Fig. S16:** Apparatus reflectively and tracking. The arena and lattice maze were both designed and colored with optimal tracking as a priority. We did not observe the mazes being tracked or reflections of the headstage LED in the mazes being tracked at any point. **a** Frames from footage recorded using our overhead camera (Fig. 1) showing the headstage LEDs pointing at the side of the lattice (left) pointing along the top bars of the lattice close to the camera (middle) or laying on a section of the arena walls (right). White lines end at tracked pixels which in all examples correspond to the 4 tracking LEDs on the headstage. **b** Color photographs showing how the tracking LEDs appear to the naked eye. **c** Frames from footage taken during recording sessions. Left) an animal faces into one corner of the area very close to the walls, reflections can be seen but these are not tracked. Inset shows a close-up of the animal’s head position with the brightness increased to highlight the reflections. Right) An animal explores the lattice maze and has his head close to the bars near one side of the maze. Again, in this case no reflections are observed or tracked.

*Tilted data rotation*

For some firing rate map analyses we computed values for the A, B and C axes of the tilted lattice. In these cases we un-tilted the tilted lattice firing rate map to align it with horizontal/vertical axes (54.735° clockwise around the X-axis and 45° clockwise around the Z-axis) using nearest neighbor interpolation (Matlab functions *affine3d* and *imwarp*), and then used the same analyses as for the aligned lattice. A similar approach was also applied to position data where necessary (Matlab functions *transformPointsForward* and *worldToIntrinsic*) using the transformation defined above or using appropriate rotation matrices (Matlab function *AxelRot*, Jacobson, M.).

*Behavior and spherical heat maps*

Using smoothed and interpolated 3D reconstructed position data we calculated the instantaneous three-dimensional heading of the animal as the normalized change in position:

$$\hat{u} = \frac{\vec{u}}{||\vec{u}||}$$

where;

$$\vec{u} = (\Delta_X(t), \Delta_Y(t), \Delta_Z(t))$$

and;

$$||\vec{u}|| = \sqrt{\Delta_X(t)^2 + \Delta_Y(t)^2 + \Delta_Z(t)^2}$$

this gives a unit vector representing the animal's heading at time *t*. We then projected these vectors on to a unit sphere and extracted position data falling within regions on the surface of the sphere corresponding to the intersection of the sphere and the 6 axes of interest: the Cartesian X (Pitch = 0°, Azimuth = 0° & 180°), Y (Pitch= 0°, Azimuth = 90° & 270°), Z (Pitch,= ±180°, Azimuth = 0°) axes and the diagonal lattice maze relative A (Pitch = ±35.26°,

Azimuth = -60° & 120°), B (Pitch = ±35.26°, Azimuth = 60° & -120°) and C (Pitch = ±35.26°, Azimuth = -180° & 0°) axes. The regions were equivalent to 60° conic sections centered on each respective axis in one direction from the origin. For each axis we combined the two corresponding directional regions. The length of time spent moving along each of these axes was calculated as the number of position samples falling along each of these axes multiplied by the sampling rate of the system. For this analysis we used only data when the animal was travelling at a speed >20cm/s. The average speed of movement along each axis was calculated separately as the total distance travelled in each axis divided by the total session time.

We also calculated the kernel smoothed density estimate of these spherical points using a Von Mises–Fisher distribution. Briefly, the Gaussian used was defined as:

$$g(x) = e^{\left(-0.5\left(\frac{x}{\sigma}\right)^2\right)}$$

where $x$ was defined as the inverse cosine of the inner dot product between each vector point and points across a sphere's surface (Matlab function *sphere*) and $\sigma$ was the standard deviation of the Gaussian, which was set to 10. In this way, the resulting three-dimensional heat plots give a density estimate of points on the sphere, where density is estimated as the sum of the Gaussian weighted distances (along the surface of the sphere) to every data point. These processes can be seen in Supplementary Video S3.

For a measure of three-dimensional thigmotaxis we calculated the total length of time spent in the inner and outer half volumes of the lattice – the inner half being an approximately 77×77×77cm cube centered on the center of the lattice. For the diagonal lattice, this central cube was rotated to match the geometry of the maze. Dwell time ratio was calculated as the ratio of these two values for each session. For the square arena environment, we omitted the Z-dimension, instead the inner half was defined as the square region with half the surface area of the arena centered on the middle of the maze.

To determine if the rats displayed a bias in their occupancy of 3D space, we also calculated the length of time the rats spent in the top and bottom of the two lattice mazes. For this we categorized data as either above the center node of the lattice (top) or below it (bottom). Dwell time ratio was calculated as the ratio of these two values for each session. As a measure of exploration coverage, for each session we calculated the proportion of lattice maze nodes (climbing bar intersections) enclosed by the convex envelope of that session's position data.

*Cluster cutting*

Single unit activity was analyzed offline using a combination of Matlab functions and custom spike sorting software. First, the dimensionality of the waveform information was reduced to the first three principal components and amplitude. Based on these parameters, an automated spike sorting algorithm (Klustakwik v3.0, [6]) was used to distinguish and isolate separate clusters. The clusters were then further checked and refined manually using a manual cluster cutting GUI (TINT v4.4.12, Axona, UK). As well as the previously mentioned features, manual cluster cutting also made use of spike auto- and cross-correlograms.

Cluster quality was operationalized by calculating isolation distance (Iso-D), $L_{ratio}$, signal to noise ratio (S/N), refractory period contamination and peak waveform amplitude, taken as the highest amplitude reached by the four mean cluster waveforms. For cluster *C*, containing $n_c$ spikes, Iso-D is defined as the squared Mahalanobis distance of the $n_c$-th closest non-*c* spike to the center of *C*. The squared Mahalanobis distance was calculated as:

$$D_{i,C}^2 = (x_i - \mu_C)^T - \sum_c^1 (xi - \mu C)$$

where $x_i$ is the vector containing features for spike *i*, and $\mu_c$ is the mean feature vector for cluster *C*. A higher value indicates better isolation from non-cluster spikes [7]. The L quantity was defined as:

$$L(c) = \sum_{i \notin C} 1 - CDF_{x_{df}^2}(D_{i,C}^2)$$

where $i \notin C$ is the set of spikes which are not members of the cluster and $CDF_{x_{df}^2}$ is the cumulative distribution function of the distribution with 8 degrees of freedom. The cluster quality measure, $L_{ratio}$ was thus defined as $L$ divided by the total number of spikes in the cluster [8]. As the signal and noise are both measured across the same impedance, signal to noise ratio (S/N) was defined as:

$$Signal\ to\ Noise\ Ratio = \left(\frac{RMS_{signal}}{RMS_{noise}}\right)^2$$

where $RMS$ is the root mean squared amplitude (maximum of the mean waveform). For $RMS_{noise}$ we used the noise cluster which accompanied unit spikes on that tetrode. This is a conservative noise measure as the noise cluster contains only noise with a high enough amplitude to breach the threshold used during recording rather than the true background noise amplitude. All four quality measures were assessed for their potential impact on our analyses by assessing the relationship between these measures and our main experimental statistics (as suggested by [7]).

*Field detection*

Unless otherwise stated, all analyses were performed on the unsmoothed firing rate maps. When detecting place fields, we looked for areas of more than 64 contiguous voxels (voxels were 50×50×50mm) with a firing rate > 20% of the maximum value. Contiguity was defined as a three-dimensional 18-connected neighborhood, which includes all voxels sharing an edge or face but not only a vertex. This was calculated using the Matlab function *bwlabeln*. Once connected regions were identified we applied two analyses: we first found their convex hull (Matlab function *convhulln*), and second, we extracted the main features of each field using the Matlab function *regionprops3*. These main features are discussed in more detail below. To be analyzed further a place field had to be visited more than 5 times, where a visit was defined as 1 second of contiguous time spent within the place field convex hull. The position of each place field was defined as its centroid: the average position of all in-field voxels. Although the open-field arena was flat, animals were free to move vertically in the area (i.e. crouching, standing and rearing) and so we treated the arena data as a thin volume; nevertheless, the analyses for the z dimension (height) in the arena must be treated with caution.

*Field volume and density*

We calculated the volume of place fields as being the sum of all its voxels. We also extracted the enclosing diameter of each place field, which was defined as the diameter of the smallest sphere capable of enclosing this convex polygon.

To quantify the relative volume of place fields and the density of fields per $m^3$ in each environment we estimated the practical volume of the lattice mazes and arena. For each recording session we calculated the volume of the convex hull (Matlab *convhulln*) of that session's position data. The practical volume of each maze was the average of these values (arena, aligned and tilted mean & SD = 0.45 & 0.12$m^3$, 1.45 & 0.16$m^3$ and 1.63 & 0.26$m^3$ respectively). Field density (fields per $m^3$) was then calculated as the number of fields expressed by each cell divided by the volume of the maze.

*Field orientation and size*

We extracted the Cartesian height and width of the fields, defined as the side length of the field region when projected on to the X, Y and Z-axes respectively (i.e. the side lengths of the minimum cuboid that could enclose the field). To examine these lengths in more detail we compared the shape of their distributions (Fig. 6d and Fig. S7). The distribution of field heights (side length along Z) was weakly bimodal in the aligned lattice, to test this we used the bootstrapped modality test described by [9] (Matlab function *bootmode*;

A. C. Penn), similar results were also found using Akaike's Information Criterion (AIC, Matlab function *fitgmdist*).

We extracted the field's orientation and principal axes, which were defined as the orientation and major axes of an ellipsoid with the same normalized second central moments as the field region. In more detail, we calculated the second central moments or covariance matrix which best described a thresholded place field, in effect, fitting a multivariate normal distribution to the field. The direction and magnitude of the best fit ellipse which describes the place field are then given by the eigenvectors and eigenvalues of this covariance matrix respectively (Fig. S17; Matlab function *regionprops3*).

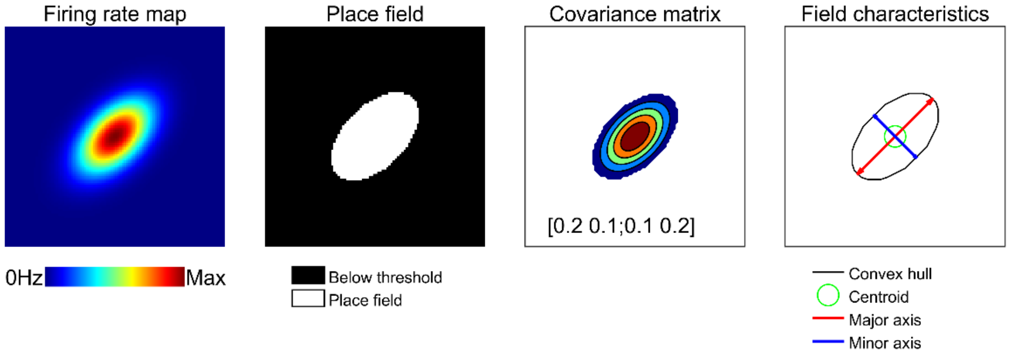


**Fig. S17:** Schematic demonstrating place field feature extraction. These examples are two-dimensional but the same principles were applied to three dimensional firing rate maps in our data. A firing rate map (left; in this case a simulated 2D Gaussian distribution) is thresholded at 20% of the peak firing rate to generate a binary field map (middle left). The second central moments of the field pixel locations give the covariance matrix which describes the best fit multivariate normal distribution of the field (middle right, visualized as a contour plot). The eigenvectors and eigenvalues of this covariance matrix give the direction and length of the place field axes. The average pixel location gives the field centroid and the convex hull of the thresholded ratemap gives the field perimeter.

We calculated the elongation index of the principal axes as:

$$Elongation = \frac{P1}{0.5(P2 + P3)}$$

where $P1$, $P2$ and $P3$ are the principal axes from largest to smallest respectively. This gives a measure of the curvature of the place field; large elongation values represent elongated fields while a value of 1 would represent a sphere. In the case of the arena, elongation was calculated using the first two largest principal axes (P1/P2). As an additional geometric measure, independent of the axis lengths, we also calculated the sphericity of each place field's convex hull, defined as:

$$sphericity = \frac{\pi^{\frac{1}{3}}(6V)^{\frac{2}{3}}}{A}$$

where $V$ is the volume of the place field and $A$ is its surface area. A sphericity of 1 would represent a sphere and any deviation from a sphere would result in a value lower than this.

To determine if fields were oriented in three-dimensions along one or more arbitrary axes we projected the place field eigenvectors (and their antipodal equivalents) onto a unit sphere. Using a similar analysis as the one described for position data, we then extracted the number of fields falling within regions on the surface of the sphere corresponding to the intersection of the sphere and the axes of interest; the Cartesian XYZ axes and the lattice maze relative ABC axes. These regions are equivalent to ~60° conic sections centered on each respective axis in one direction from the origin, thus for each axis we combined the two corresponding directional regions.

To determine whether the number of fields oriented parallel to one axis was greater or less than another axis we calculated 95% confidence intervals for each point. To do this we extracted the orientations of random place fields with replacement so that the number of random field orientations was equal to the number of observed fields. We then recalculated the proportion of fields in this shuffle that were oriented parallel to each axis as above. We did this 1000 times for each maze. The error bars shown in Fig. 7c represent the $2.5^{th}$ and $97.5^{th}$ percentile ranks of these shuffled counts. If the observed value for one axis fell within the error bars of another axis the two axes were not considered to differ significantly.

To determine whether more fields were parallel to an axis than would be expected by chance, we generated 1000 random points on the face of a sphere and counted the proportion of points falling within the area around each axis. We did this 1000 times. Chance was calculated as the interval between the 2.5$^{th}$ and 97.5$^{th}$ percentile ranks of this distribution. If the observed field count for an axis exceeded the upper threshold it was considered to be overrepresented with respect to chance.

As fields were observed to fall in alignment with the XYZ axes in the aligned lattice maze and the ABC axes in the diagonal lattice we also computed an 'axis ratio' for comparison. This was defined as:

$$axis\ ratio = \sum XYZ / \sum ABC$$

To test the likelihood of observing these axis ratios by chance we compared them to a shuffled distribution. For this, we randomly distributed 1000 points across the surface of a sphere 1000 times and recomputed the above values. If the axis ratio of a maze exceeded the 1$^{st}$ or 99$^{th}$ percentile of the ratios obtained in the shuffle it was defined as significantly deviating from 1 (no axis bias of any kind). In this case an observed value less than the 1$^{st}$ percentile would represent a significant overrepresentation of fields aligned to the ABC axes, while a value exceeding the 99$^{th}$ percentile would represent a significant overrepresentation of fields aligned to the XYZ axes.

For visualization, we calculated the Von Mises–Fisher kernel smoothed density estimate of these place field vectors across the sphere's surface (as described in Behavioral analyses). These 3D spherical maps are presented in the main text for visualization only (Fig. 2&7).

In addition to the place field orientation analyses presented in the main text, we also generated spherical field maps predicting the pattern of results in each maze if all fields were parallel to the maze axes. These maps were produced by normalizing the output of our Von Mises–Fisher kernel smoothed density estimate computed on the directional angles associated with each maze axis (i.e. the intersection points of each maze axis with a unit sphere). In the case of the open field arena we included only the X and Y axes, for the aligned lattice we included the X, Y and Z axes and for the tilted lattice we only included the A, B and C axes. This process can be seen in Fig. S8a. These maps essentially highlight the pitch × azimuth regions associated with the maze axes.

Next, we correlated the 2D cylindrically projected maps observed in each maze (i.e. the actual, collected data) with each of these possible predictive maps (Pearson, pairwise correlation, Matlab function *corr*) also projected cylindrically. In this way, if the fields in a maze are parallel to the maze axes we would expect a high correlation between the observed and predicted maps. To test if these correlations were significantly higher than would be expected by chance (and to account for the loss of sphericity in these cylindrical projections) we generated distributions of shuffled maps for each maze. For these shuffles we generated 1000 maps as above using 10000 random spherical points (an example can be seen in Fig. S8a). We correlated the shuffle maps with the observed data and extracted the $99^{th}$ percentile of this distribution. The original correlations were deemed statistically significant if they exceeded this value.

*Field elongation and sphericity*

For each place field we tested whether its elongation index and sphericity deviated significantly from a distribution that would be expected by chance using an analysis inspired by one reported previously [10]. For each place field we defined a perfect sphere, centered on the field's centroid. The diameter of this sphere was calculated such that it would share the same convex volume (the volume of the convex hull enclosing the field voxels) as the place field. This was calculated as:

$$equivalent\ diameter = 6(\frac{vf}{\pi})^{\frac{1}{3}}$$

where *vf* is the convex volume of the place field. We found this to be more accurate than the geometric mean approach reported previously [10] which assumes all place fields are perfectly elliptical and thus tends to underestimate equivalent diameter. Next, the spikes emitted within the place field were randomly shuffled among the trajectories through this sphere using a multivariate Gaussian process (Matlab *normrnd*). The mean of the Gaussian was the sphere center and the standard deviation of the Gaussian was set to 1.8 × the radius of the sphere (to approximately match the 20% thresholding used during field detection). Each spike was then assigned to the position of the nearest trajectory data point (Matlab *knnsearch*). The result of this procedure was a normally distributed point cloud of spikes centered on the centroid of the original field with the same equivalent diameter and firing rate.

We then recomputed the firing rate map for these shuffled spikes and extracted its elongation index as described above. This procedure was repeated 100 times for each place-field. Place fields with an elongation index or sphericity that could be expected, by chance, from an underlying spherical field (i.e. with an elongation index lower than the 95$^{th}$ percentile rank of the shuffled distribution) were defined as spherical or isotropic: otherwise, place fields were defined as non-spherical or anisotropic.

*Field distribution*

To test whether place fields in the mazes were distributed homogenously we compared the distribution of place field centroids in X, Y and Z to a shuffled set of centroids. We excluded place fields falling outside the lattice frame to limit the test to a consistent and repeatable volume. We then calculated the median position of fields in the X, Y and Z axes. Next we generated *N* uniformly random points (Matlab function *rand*) within the lattice frame, where *N* was the number of real place fields falling within this volume. We repeated this

1000 times and at each step calculated the median position of the centroids in X, Y and Z. Next we calculated the 2.5th and 97.5th percentile ranks of these distributions; if the observed median position in X, Y or Z was found to exceed these bounds we considered that place fields were not distributed uniformly around the center of the maze in that axis.

*Binary morphology*

We also conducted a binary morphological analysis on thresholded firing rate maps to detect their maximal connectivity along each dimension (Fig. S18). First, binary firing rate maps (*A*) were generated by thresholding firing rate maps (*F*) at 10% of their maximum value:

$$F(x) = \begin{cases} 1 & if\ x \geq 0.1(\max(F)) \\ 0 & if\ x < 0.1(\max(F)) \end{cases}$$

We then performed morphological erosion (*Θ*) using structuring element vectors (*B*) with lengths ranging from 3-19 voxels along each cardinal axis (Matlab functions *imbinarize* and *bwhitmiss*):

$$A\ \theta\ B = \{z \in E \mid B_z \subseteq A\}$$

For each erosion we took the linear sum of the remaining voxels as a measure of the map's connectivity along that dimension and then expressed this as the proportion of all remaining voxels for that element length. This last step was necessary to account for the fact that as the structuring element increases in size the likelihood of voxel connectivity decreases substantially, although we achieved similar results without it.

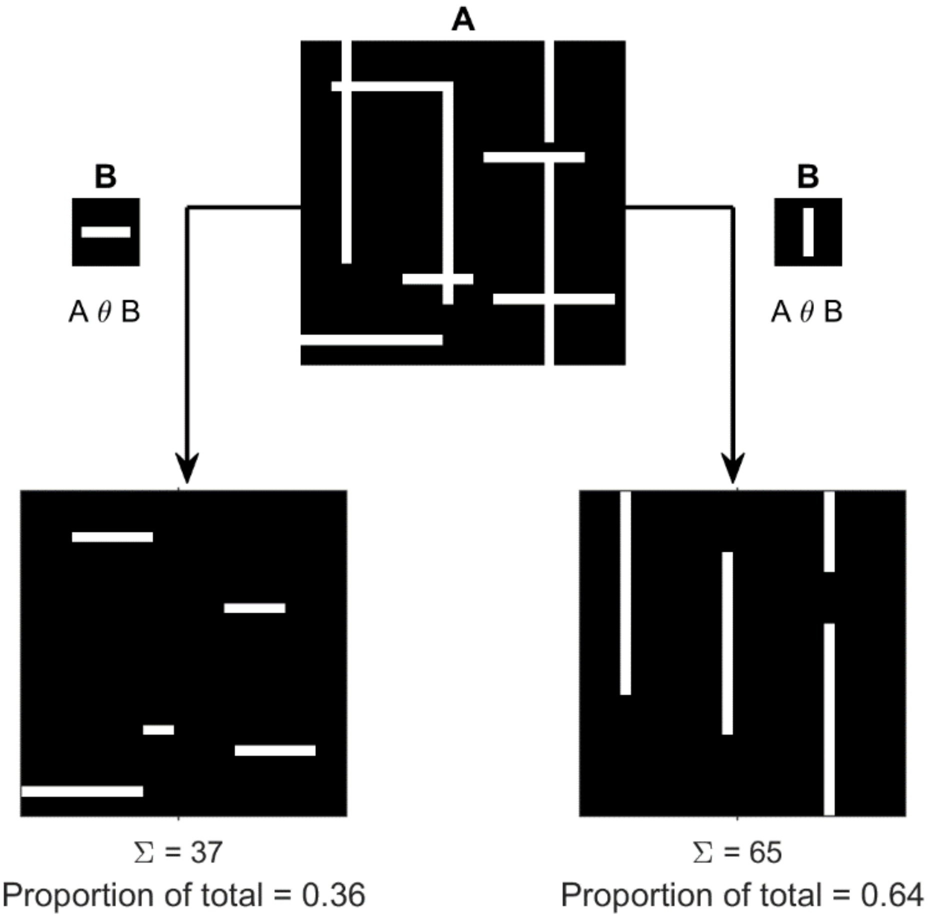


**Fig. S18:** Two dimensional representation of the binary morphological process. A firing rate map is thresholded to form a binary image (A), where 1's represent voxels with a value above 10% of the map's maximum and 0's represent all other voxels. This binary map is then eroded using structuring elements (B) which are vectors of varying lengths that are parallel to one of the primary cardinal exes. The result is a map where 1's represent voxels that can accommodate the structuring element and 0's are voxels that cannot. As a measure of relative connectivity along each dimension, we took the sum of each eroded map and expressed this as a proportion of the sum of all maps. In this example, A has longer vertical periods of connectivity than horizontal ones, this is confirmed by the fact that almost two thirds of the eroded voxels can be found after eroding with a structuring element parallel to the Y-axis.

*Autocorrelation and spatial information*

Anisotropic place fields that are oriented parallel to a Cartesian axis (i.e. forming vertical columns or horizontal bands) will be visible on multiple two-dimensional slices through a three-dimensional rate map (Fig. S9). To investigate this possibility we computed the three dimensional autocorrelation, $r$, of each place cell's firing rate map, defined as:

$$r(\tau_x, \tau_y, \tau_z) = \frac{M \sum_{x,y,z} \lambda(x,y,z)\lambda(x-\tau_x, y-\tau_y, z-\tau_z) - \sum_{x,y,z} \lambda(x,y,z) \sum_{x,y,z} \lambda(x-\tau_x, y-\tau_y, z-\tau_z)}{\sqrt{\left[M \sum_{x,y,z} \lambda(x,y,z)^2 - \left[\sum_{x,y,z} \lambda(x,y,z)\right]^2\right] \left[M \sum_{x,y,z} \lambda\left(x-\tau_x, y-\tau_y, z-\tau_z\right)^2 - \left[\lambda(x-\tau_x, y-\tau_y, z-\tau_z)\right]^2\right]}}$$

where $\lambda(x,y,z)$ is the firing rate at the location $(x,y,z)$ in the firing rate map, $M$ is the total number of voxels in the rate map, and $\tau_x$, $\tau_y$ and $\tau_z$ correspond to $x$, $y$, and $z$ coordinate spatial lags [11]. From this, we extracted the three voxel wide midline portion along the X, Y and Z axes and took the median value of the all the contained correlation scores. We also extracted the values falling exactly on these midlines (Matlab function *interp3*) for a measure of similarity over increasing distances or autocorrelation voxel lag (see Fig. S19 for a schematic and examples).

We also projected firing rate maps onto the three possible Cartesian planes by taking the average along each axis (ignoring empty voxels). We then calculated the spatial information content found in each projection and expressed these as the proportion of total spatial information. This last step was to account for cells with different overall spatial information content.

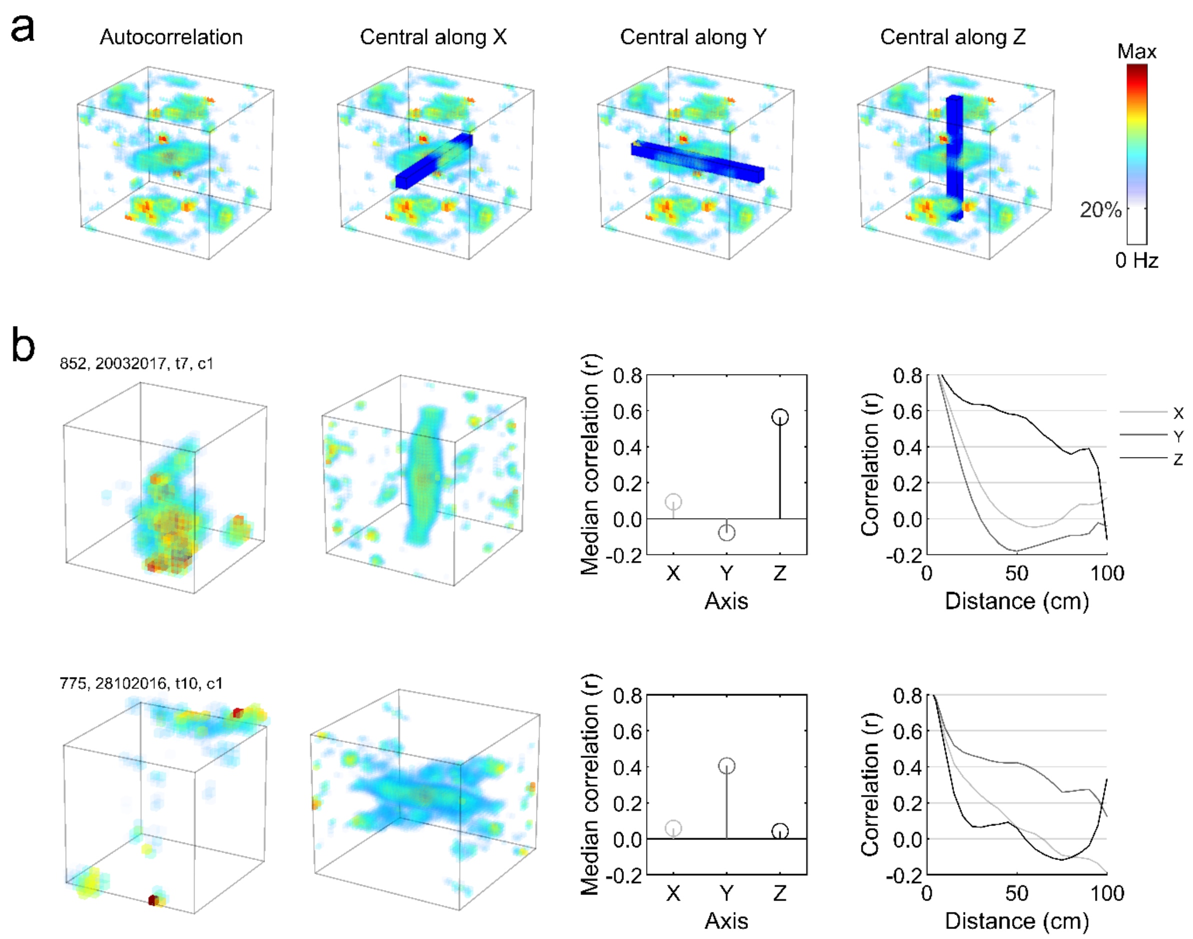


**Fig. S19:** Example autocorrelations and schematic of the autocorrelation analysis procedure. To quantify the self-similarity of firing rate maps along each maze axis we generated firing rate map autocorrelations using the method described above. An example of one of these can be seen in **a**. Next we extracted the values falling along the midline of each axis (blue regions in **a**). For each of these we calculated the overall median correlation value. We also extracted values falling on the midline of each axis (i.e. a line running through the middle of each blue region). **b** The left column shows two example firing rate maps (one per row), the second column shows the result of an autocorrelation performed on each rate map. The third column shows the median autocorrelation value calculated for each axis (blue regions in **a**). The right column shows the values found along the midline of each axis of the autocorrelation; as the autocorrelations are symmetrical we have only shown the positive half of each axis. Note that for the top cell with vertically aligned firing the Z-axis median value is highest and this axis exhibits generally higher correlation values along this axis. The bottom cell has horizontally aligned firing and as a result the same effect is instead exhibited by the Y-axis.

*Place field standard deviation analysis*

To quantify the standard deviation of firing in the X, Y and Z dimensions for each place field, we summed the field along each of these dimensions and fitted a Gaussian (Matlab function *fit*, with curve *gauss1*) to the result. We then extracted the standard deviation of this Gaussian as a measure of place field spatial variance. Values resulting from poor Gaussian fits ($r^2 < 0.2$) were excluded.

*Trajectory downsampling*

To test if spatial information and autocorrelation were affected by the inhomogeneous sampling of space in the aligned lattice (i.e. animals move vertically much less than horizontally) we used a downsampling procedure. This consisted of filtering trajectory and spike data to include 50% vertical movements (defined as movements at a pitch >30° or <-30°) and 50% horizontal movements (defined as movements at a pitch <30° and >-30°) for a total trajectory made up of sub-sampled but homogeneous data. We used a pitch angle of 30° to delineate movements as this divides the face of a sphere into two halves with equal surface area (a 'belt' around the equator for one half and two spherical caps for the other). We generated firing rate maps using this sub-sampled trajectory and spike data and then repeated the analyses described in Supp. Methods: *Autocorrelation and spatial information*. We repeated the same test using data subsampled to include 50% inner (inner 50% volume of the lattice) and outer (outermost 50% volume) movements or 50% top (top 50% volume of the lattice) and bottom (bottom 50% volume) movements.

*Zingg shape categorisation*

To quantify the shape of place fields we used the method described by Zingg [12]. If P1, P2 and P3 are the long, intermediate, and short axes of a place field and R is a number greater than one then four mutually exclusive shape classes can be defined (see Table S1). In our approach we used R = 3/2 as suggested by Zingg [12]. This approach classifies shapes into four mutually exclusive groups that can be used to evaluate the shape of individual fields; 'equant', 'prolate', 'oblate' and 'bladed'. Of greatest interest to us are the equant (spherical) and prolate (significantly elongated) classes. For each maze we calculated a shuffled distribution for comparison. To maintain a suitable underlying distribution we randomly shuffled the principal axis lengths of our place fields, grouped them into triplets and sorted them from highest to lowest. We then recalculated the number of triplets falling into each shape class. We repeated this process 1000 times. If the number of fields fulfilling a shape class in a given environment exceeded the 99$^{th}$ percentile of the number of fields fulfilling the same class in the shuffles this was considered a significant deviation from chance.

**Table S2**

*Table showing Zingg (1935) shape classification criteria*

| Shape class | P1 and P2 | P2 and P3 | Description | Example |
|---|---|---|---|---|
| Equant | P2 < P1 < R*P1 | P3 < P2 < R*P3 | all dimensions are comparable | Sphere |
| Prolate | P1 > R*P2 | P3 < P2 < R*P3 | one dimension is much longer | Cigar |
| Oblate | P2 < P1 < R*P2 | P2 > R*P3 | one dimension is much shorter | Pancake |
| Bladed | P1 > R*P2 | P2 > R*P3 | all dimensions are very different | Sheet of paper |

*Comparing activity between mazes*

We sought to compare some basic firing properties between mazes and determine if there was a link between the characteristics of fields in the arena and lattice mazes. Overall maze firing rates were calculated as the total spikes emitted in a maze divided by the total time spent there. Spatial information content was calculated as in Methods: *Place cell criteria*. Sparsity was defined as:

$$sparisty = \sum \left(P_i R_i^2\right)/R^2$$

where $P_i$ is the probability of occupancy of bin $i$, $R_i$ is the mean firing rate in bin $i$, and R is the overall mean firing rate [13]. To compare field elongation between mazes we correlated the elongation of fields in the arena (or average elongation if a cell expressed multiple fields) and the elongation of fields (or average) in the following lattice maze session.

To investigate if multiple fields of the same cell shared similar characteristics we compared their length and orientation, these analyses were conducted only on those cells with more than 1 place field. For length we calculated the average pairwise difference in length between the place fields of a cell and compared this to a shuffled distribution containing the average pairwise differences between randomly paired fields selected from the overall dataset (for that maze) without replacement. For orientation we calculated the average pairwise inner angle between the major axes of place fields and compared this to a shuffled distribution containing the average pairwise angles between randomly paired fields selected from the overall dataset (for that maze) without replacement. Lastly, to determine if cells were likely to exhibit the same number of fields in each maze, we calculated the difference between total fields in the lattice mazes and arena. We compared these values to shuffles where we calculated the same difference between randomly paired cells selected from the overall dataset (for that maze) without replacement.

*Local field potential (LFP) analyses*

Before analysis, all LFP data were removed of their direct current offsets, slowly changing components, and running line noise using the Chronux toolbox [14] *locdetrend* function which subtracts the linear regression line fit within a 1s moving window. They were then resampled at 250 Hz using a polyphase anti-aliasing filter (MATLAB function *resample*, pchip interpolation).

To obtain a theta phase angle for each spike, LFPs were first bandpass filtered in the 6-12 Hz range (fourth-order Butterworth, Matlab functions *butter* and *filtfilt*) before a Hilbert transform was applied to obtain the instantaneous phase angle (Matlab function *hilbert*). Instantaneous frequency was calculated as the derivative of this analytic signal (Matlab function *instfreq*) and instantaneous amplitude was calculated as its magnitude.

To assess the relationship between running speed and the theta oscillation we compared the instantaneous theta power/amplitude at every position data point (every 20 ms) to the animals' instantaneous running speed. Instantaneous speed was estimated as the total distance travelled in every 40 ms window (i.e. the distance between every position data sample, the previous one and the next one divided by the time between them). To quantify the relationship between speed and power we fitted a linear regression model using a least-squares approach (Matlab function *polyfit*, 1 degree) and extracted the slope, y-intercept and sum of squared error. To assess the curvature of these relationships we also fitted a power function ($y(x) = ax^b + c, where\ a \geq 0$) to each session's speed-power curve and extracted the power parameter $b$ (Matlab function *fit*). In this model $b$=1 denotes the linear function $f(x) = x$, $b<1$ denotes a downward curve and $b>1$ denotes an upward curve. We also performed the same procedures to test the relationship between running speed and instantaneous frequency.

To calculate global/overall theta characteristics we computed average power spectral densities (PSDs) for each recording session by first zero-padding data to the next highest power of 2. A Welch spectral estimator was then applied to obtain the PSD (Matlab function *pwelch*, Hamming window, 8 segments, 50% overlap). This was computed for 500 logarithmically spaced points between 0-250 Hz. Theta power was estimated as the maximum power found in the theta band (6-12Hz), theta frequency was defined as the frequency associated with this maximum power.

*Running speed analyses*

Instantaneous running speed was estimated as the total distance travelled in every 40ms window. For each cell, instantaneous firing rate was estimated as the smoothed spike histogram (20 ms bins, 13 bin or 260 ms Gaussian smoothing window using Matlab function *fspecial*). To quantify the relationship between speed and firing rate we used an analysis similar to that described previously [15]. We binned the animals' running speeds in 2cm/s increments and calculated the mean firing rate for each running speed bin and the total time spent moving at that speed. We then fitted a linear regression model to the average firing rate/speed data using a least-squares approach (Matlab function *polyfit*, 1 degree) and extracted the slope, y-intercept and sum of squared error.

*Spike phase and autocorrelation analyses*

To quantify the intrinsic theta modulation of every place cell we used an analysis described previously [16,17]. For each cell we calculated the ±500ms spike autocorrelation in 10ms bins, normalized this to the maximum value found between 100 and 150 ms and removed values >1. Then we fit the following function to the remaining data:

$$y(t) = \left(a * \left(\sin\left(2\pi\omega t + \frac{\pi}{2}\right) + 1\right) + b\right) * \exp\left(-\frac{|t|}{\tau_1}\right) + c * \exp\left(-\frac{r^2}{\tau_2^2}\right)$$

where $a, b, c, \omega, \tau_1$ and $\tau_2$ were fit to the data using a non-linear least squares method (Matlab function *fit*) and $t$ is the autocorrelogram time lag. In simple terms this function fits a sine wave of frequency $\omega$ to the data and the exponential term allows for this to decrease exponentially as the time lag increases (reflecting the exponential decay inherent to all spike autocorrelations). The last Gaussian term helps to center the fit on the autocorrelogram peak, which we found to be unnecessary in most cases. A measure of theta modulation strength was defined as $a/b$, which intuitively corresponds to the ratio of the sine fit relative to the baseline in the autocorrelogram. The parameter $\omega$ was extracted as the intrinsic theta modulation of the cell. We restricted possible values for $\omega$ to [6, 12], $a$ and $b$ were restricted to non-negative values [0 Inf], $c$ was restricted to [0, 0.8], $\tau_1$ was unrestricted and $\tau_2$ was restricted to [0, 0.05]. This fitting procedure was only carried out on cells that fired at least 500 spikes.

For each cell the instantaneous theta phase of every spike was calculated by linear interpolation of the instantaneous theta phase signal described previously. These phase angles were binned between $-\pi$ and $\pi$ in 0.1 rad bins. The cell's preferred theta phase was defined as the circular mean of these angles and the strength of this modulation was defined as the mean resultant vector length of these angles (Matlab functions *circ_mean* and *circ_r* respectively, circular statistics toolbox, [18]). At the population level, all cell preferred phases were collated, binned between $-\pi$ and $\pi$ in 0.1 rad bins and again we calculated the strength of this modulation using the mean resultant vector length as above. Distributions were compared between mazes using a two-sample Kuiper test (Matlab function *circ_kuipertest*, circular statistics toolbox, [18]) and the phase lag between them was estimated by cross-correlation (Matlab function *finddelay*).

*Histology*

At the end of the experiment animals were given an overdose of pentobarbital intraperitoneally (Euthatal, Merial Animal Health Ltd., Essex, UK), and perfused with 0.9%

saline solution followed by a 4% formalin solution. The brain was extracted and stored in 4% formalin for at least seven days prior to any histological analyses. The brains were sliced in 30 μm sections on a freezing microtome at −20°. These sections were stained with a 0.1% cresyl violet solution and the slice best representing the electrode track was then imaged and color corrected.
